# Liquid–liquid phase separation of α-synuclein is highly sensitive to sequence complexity

**DOI:** 10.1101/2023.08.03.551831

**Authors:** Anindita Mahapatra, Robert W. Newberry

**Affiliations:** Department of Chemistry, The University of Texas at Austin, 105 E 24th St. Austin, TX, 78712, USA

## Abstract

The Parkinson’s-associated protein α-synuclein (α-syn) can undergo liquid–liquid phase separation (LLPS), which typically leads to the formation of amyloid fibrils. The coincidence of LLPS and amyloid formation has complicated the identification of the molecular determinants unique to LLPS of α-syn. Moreover, the lack of strategies to selectively perturb LLPS makes it difficult to dissect the biological roles specific to α-syn LLPS, independent of fibrillation. Herein, using a combination of subtle missense mutations, we show that LLPS of α-syn is highly sensitive to its sequence complexity. In fact, we find that even a highly conservative mutation (V16I) that increases sequence complexity without perturbing physicochemical and structural properties, is sufficient to reduce LLPS by 75%; this effect can be reversed by an adjacent V-to-I mutation (V15I) that restores the original sequence complexity. A18T, a complexity-enhancing PD-associated mutation, was likewise found to reduce LLPS, implicating sequence complexity in α-syn pathogenicity. Furthermore, leveraging the differences in LLPS propensities among different α-syn variants, we demonstrate that fibrillation of α-syn does not necessarily correlate with its LLPS. In fact, we identify mutations that selectively perturb LLPS *or* fibrillation of α-syn, unlike previously studied mutations. The variants and design principles reported herein should therefore empower future studies to disentangle these two phenomena and distinguish their (patho)biological roles.

## Introduction

Liquid–liquid phase separation (LLPS) is the metastable assembly of macromolecules into liquid-like droplets, which enables spatiotemporal regulation of cellular functions (1). Fibrillation, on the other hand, is the aberrant self-assembly of functional proteins into amyloid fibrils, which is a pathological hallmark of various neurodegenerative disorders (2; 3). Recently, LLPS has been implicated as a regulator of pathological fibrillation (4; 5), primarily because several amyloid-forming proteins have been found capable of undergoing LLPS in physiological conditions (6-11), and disease-associated mutations that affect fibrillation of these proteins have been found to affect their LLPS as well (11-14). Prominent examples include hnRNPA1 (6; 7), TDP-43 (7), and FUS (8; 9) – associated with amyotrophic lateral sclerosis (15); and tau (10) – associated with Alzheimer’s disease (16). A relatively new inclusion in this group is α-synuclein (α-syn) (11) – associated with Parkinson’s disease (PD), (17; 18), which has only recently been found to undergo LLPS *in vitro* (11; 19-21), in human cell lines (11), and in *C. elegans* (22).

α-Syn is an intrinsically disordered, presynaptic protein, the fibrillation of which is implicated in the pathogenesis of PD and other synucleinopathies (17; 23). Native α-syn is involved in the trafficking of synaptic vesicles *via* membrane binding (24; 25), and why and how it converts to pathogenic amyloid fibrils is still not clearly understood. *In vitro* studies have established that the process involves structural conversion of the soluble monomer into a fibrillation-competent nucleus, which goes through oligomeric and pre-fibrillar intermediates to eventually form insoluble fibrillar aggregates (26). The rate of α-syn fibrillation is known to be influenced by environmental triggers (27), as well as various missense mutations in the amino acid sequence of α-syn that have been directly linked to the onset/progression of familial and sporadic PD (28; 29). The recent discovery of α-syn’s ability to undergo LLPS, followed by a gradual liquid-to-solid transition into amyloid-like fibrillar aggregates (11), has complicated matters further and led to significant interest in the potential role of α-syn LLPS in fibrillation. Several studies have investigated LLPS of α-syn in presence of extrinsic factors that affect its fibrillation, such as pH (30), metal ions (11; 31; 32), and natural products (33). However, the intrinsic drivers of α-syn LLPS are still largely unclear. Reports to date have implicated all three domains of the protein (*i*.*e*., N-terminal, non-amyloid-β component or NAC, and C-terminal) to be involved in mediating its LLPS, either by electrostatic or hydrophobic interactions (11; 19); but more specific, residue-level determinants of α-syn LLPS remain unknown, making it difficult to ascertain its biological significance. Moreover, the coincidence of α-syn LLPS and fibrillation complicates the dissection of their individual roles in disease.

Although α-syn is structurally and functionally distinct from other phase-separating, amyloidogenic proteins like hnRNPA1, TDP-43, FUS and tau, they all share at least one notable similarity: the presence of low-complexity domains (LCDs) in their primary structure. LCDs are segments within proteins that have low compositional (or sequence) complexity, *i*.*e*., they are composed of relatively few distinct amino acids. LCDs are widely implicated as the key drivers of protein phase separation. They have been found sufficient for LLPS of hnRNPA1 (7), TDP-43 (13), and FUS (34); and are generally believed to mediate LLPS through transient multivalent interactions between specific types of amino acids that are prevalent within them, mostly aromatic and/or polar residues (4; 35; 36). For example, LLPS of FUS is mediated by tyrosine and glutamine residues in its LCD (37; 38), and LLPS of TDP-43 is mediated by arginine and regularly spaced aromatic residues in its LCD (39-41). The role of LCDs in protein LLPS has therefore largely been ascribed to the physicochemical properties, and often identities, of their constituent amino acids (4). However, given the diversity of composition between different LCDs, we wondered if their complexity *itself* might be a key determinant of LLPS, and sought to test this hypothesis in α-syn, which is predicted to contain two LCDs (11).

Some previous studies examined the roles of LCDs in protein LLPS by deleting parts or whole of the LCDs from the respective proteins (6; 40); although informative, this approach can significantly alter one or more fundamental physicochemical properties of the protein, including its size, hydrophobicity, and charge. Because we wanted to specifically investigate the impact of sequence complexity on α-syn LLPS, we took a different experimental approach that would largely preserve the fundamental physicochemical properties of the protein while only modifying the complexity of its two LCDs. To achieve this, we employed the Simple Modular Architecture Research Tool (SMART) – a computational tool that engages the SEG algorithm to identify LCDs in protein sequences (42). The SEG algorithm (43-45) classifies a protein segment as LCD based solely on its compositional complexity, which is determined by the count of unique amino acids in the given segment, independent of sequence patterns or repetition (45). Mathematically, compositional complexity of a protein segment of length *L* is defined by 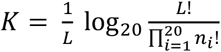, where *n*_*i*_ is the number of occurrences of each of the 20 different amino acids in the given segment, with 0≤*n*_*i*_≤*L* and 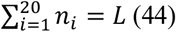. For the purpose of our study, we defined sequence complexity of α-syn as the cumulative compositional complexity of the two segments corresponding to its two LCDs; and by careful design, introduced chemically subtle missense mutations in one or both of these segments, so that only sequence complexity of α-syn is altered without affecting its structural and physicochemical properties. By comparing the propensity of the resulting variants to phase separate relative to the WT, we demonstrated that LLPS of α-syn decreases dramatically with increasing sequence complexity. We also showed that our designed mutations have no consistent effect on α-syn fibrillation in the absence of LLPS; in doing so, we identified mutations that selectively perturb either LLPS *or* fibrillation of the protein, providing new tools for studying the behaviour of α-syn.

## Results

### Designing mutations to test the role of sequence complexity in LLPS of α-syn

α-Syn contains two low-complexity domains (LCDs) in its primary structure, as predicted by SMART (42) (Fig. 1, WT). To test the role of sequence complexity in α-syn LLPS, we designed subtle missense mutations in these two LCDs (LCD1 and LCD2), such that they increase sequence complexity of the protein while minimally perturbing its structural and physicochemical properties (see Materials and Methods for detailed explanation of our design strategy and quantification of sequence complexity). After examining the sequence of LCD1 (residues 10–23), we identified a single valine-to-isoleucine mutation (V16I) that would be sufficient to increase the sequence complexity of this region above the threshold for identification as a LCD by SMART (Fig. 1, variant –LCD1). Owing to the chemical subtlety of this mutation (a single methyl group on a single hydrophobic residue in a 140-residue protein), computational analyses predicted minimal changes to the protein’s charge and hydrophobicity (Table S1), as well as its propensity for disordered or helical structure (Figs. S1A and C). Furthermore, to test if restoring the original sequence complexity can rescue LLPS behaviour of the WT protein, we designed another variant in which the disruption of LCD1 by the V16I mutation was rescued by introducing a second V-to-I mutation (V15I) adjacent to the V16I (Fig. 1, variant LCD1*). As a result, this variant had the same sequence complexity as the WT protein, albeit having a slightly different amino acid sequence (with two isoleucine residues instead of two valine residues at positions 15 and 16). SMART analysis predicted that introducing a single V-to-I mutation in LCD2 (residues 63–78), would not sufficiently increase sequence complexity to disrupt this LCD. We therefore paired a V-to-I mutation (V70I) with a glycine-to-proline mutation (G68P), which together raised sequence complexity of this region above the threshold for detection as a LCD by SMART (Fig. 1, variant –LCD2). We also combined the –LCD1 and –LCD2 substitutions to make a variant with no predicted LCDs (Fig. 1, variant –LCD1&2). Like the –LCD1 variant, the LCD1*, –LCD2 and – LCD1&2 variants were all predicted to retain the physicochemical properties and structural propensities of the WT protein, due to the chemical subtlety of the introduced mutations (Table S1, Figs. S1A and C). We therefore expected our designed LCD variants to test the role of sequence complexity in LLPS of α-syn.

**Figure 1.**
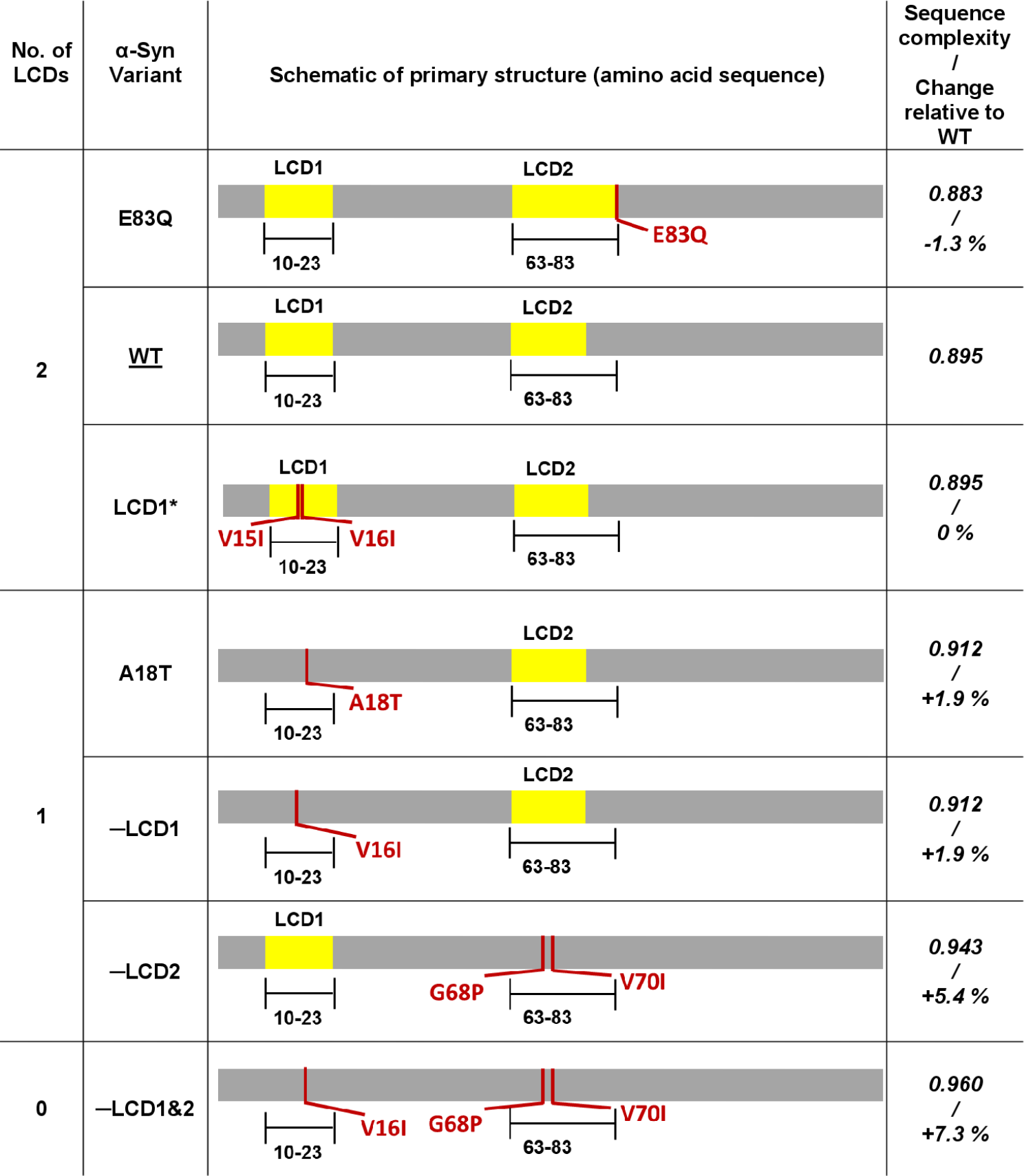
In silico analyses of amino acid sequences of α-syn variants using SMART, showing their respective LCDs and calculated sequence complexities. The calculation of sequence complexity is described in detail in ‘Quantification of sequence complexity of LLPS variants’ under the Materials and Methods section.

Curious to see if any of the known PD-associated mutations of α-syn affect any of its LCDs, we ran SMART prediction on the amino acid sequences of all known patient variants. Interestingly, only the sporadic PD-associated mutation A18T (29) (located within the LCD1 region of α-syn) was found to sufficiently increase compositional complexity of this region so that it was no longer predicted as a LCD by SMART (similar to V16I). The familial PD mutation E83Q (46) (lying slightly outside the LCD2 region of the WT protein) was predicted by SMART to slightly increase the length of this region (Fig. 1, variant E83Q *vs*. WT). Although the E83Q mutation slightly lowered sequence complexity relative to WT α-syn, it did not change the number of predicted LCDs of α-syn, and may therefore be expected to behave similar to the WT protein. Interestingly, both E83Q and A18T, are known to promote fibrillation of α-syn *in vitro* (29; 46). Though the patient variants did not have the same charge, hydrophobicity and helical propensity as the WT protein, unlike our carefully designed synthetic LCD variants, they might be useful for exploring the role of sequence complexity and/or LLPS in PD pathogenesis.

### LLPS of α-syn decreases with increasing sequence complexity

To compare the propensity of droplet formation among α-syn variants with different sequence complexities, we induced *in vitro* LLPS of all seven variants (including WT) by introducing a widely used molecular crowder (PEG 8000, 20% w/v) under physiological salt and pH conditions (PBS, pH 7). Consistent with previous observations under similar conditions (30), droplets of the WT protein started forming almost immediately (Fig. S2). However, as the nascent (0-hour) droplets were too mobile to be focused under the fluorescence microscope for imaging, LLPS samples of all variants were allowed to mature for up to 4 hours at 37 °C prior to imaging and quantification. Quantification of the area covered by droplets revealed a marked decrease in droplet formation of α-syn upon decreasing the number of LCDs by increasing sequence complexity of the protein (Fig. 2). A similar correlation between sequence complexity and phase separation was observed when droplets were quantified by turbidity (Fig. S3).

**Figure 2.**
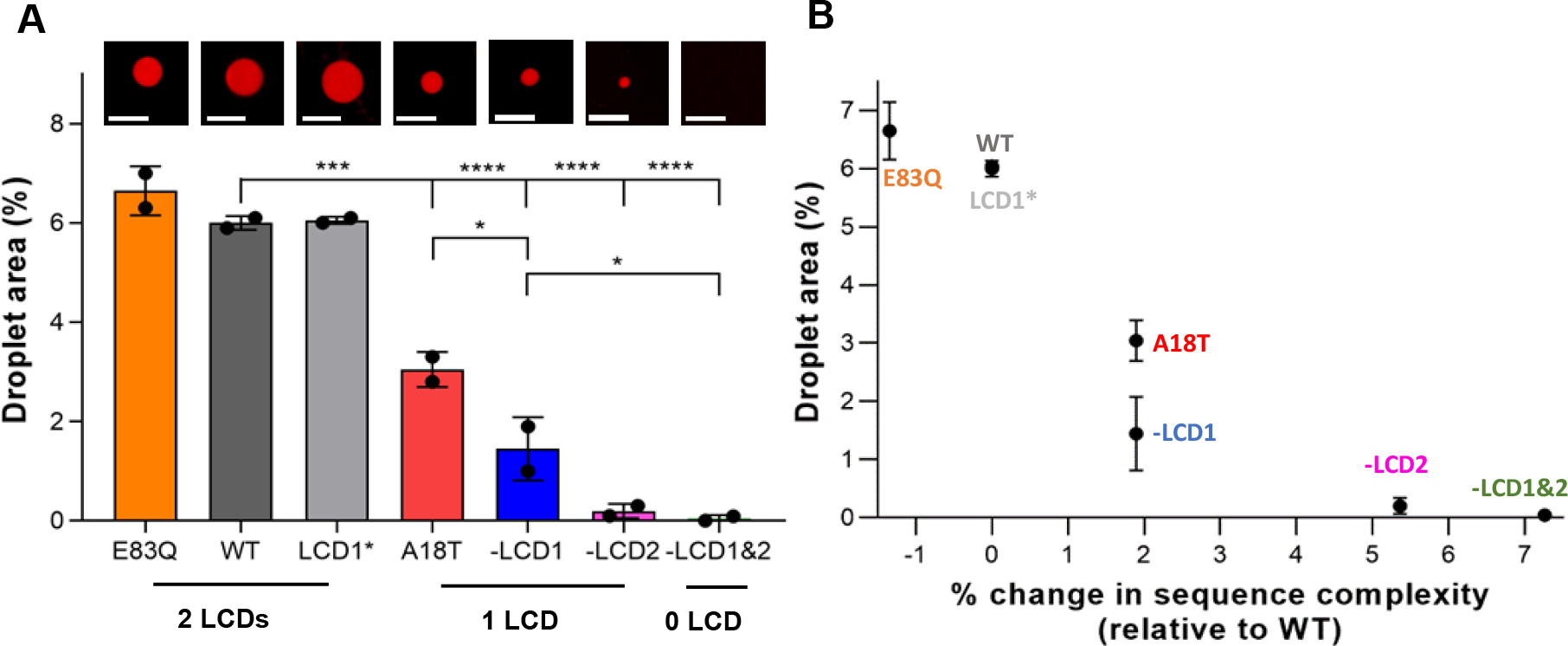
**(A)** Quantification of droplet formation for all LLPS variants of α-syn at pH 7, showing decrease in amount of droplet formation with decreasing number of LCDs in the protein. Images over each column show representative high magnification (126x) confocal microscopic images of individual droplets of respective LLPS variants. Scale bar = 10 μm. **(B)** Plot of droplet area *vs*. change in sequence complexity (relative to WT α-syn), showing a decrease in LLPS propensity with increase in sequence complexity of α-syn. Data represent mean ± standard deviation (SD) for n = 2 independent experiments. Statistical significances are indicated by *: p ≤ 0.05, **: p ≤ 0.01, ***: p ≤ 0.001, ****: p ≤ 0.0001.

Even a single V-to-I mutation in the first LCD (V16I), leading to disruption of this LCD due to increased sequence complexity, was sufficient to reduce droplet formation by a dramatic 75%, despite its minimal effects on the protein’s physicochemical and structural properties (variant –LCD1). Remarkably, the reduction in α-syn LLPS by the V16I mutation was reversed by a second adjacent V-to-I mutation (V15I), which restores the original sequence complexity and hence LCDs of the WT protein (variant LCD1*). The complete rescue of α-syn LLPS upon introducing this second V-to-I mutation shows that V-to-I mutations do not inherently decrease LLPS of α-syn; rather the overall sequence complexity of the protein appears to govern its propensity to phase separate.

Disrupting the second LCD (LCD2) of α-syn by sufficiently increasing sequence complexity through the G68P and V70I mutations, was also found to drastically reduce LLPS relative to the WT (variant –LCD2). In contrast, a mutation in the LCD2 region that does not disrupt this LCD (such as V70I) could not reduce LLPS relative to WT α-syn (Fig. S4). The drastic reduction in LLPS propensity of the –LCD2 variant is consistent with previous observations by Ray S. *et al*. indicating the importance of the NAC domain (residues 61–95, containing the LCD2 region) in facilitating α-syn LLPS (11). For the –LCD1&2 variant, with highest sequence complexity relative to the WT and no predicted LCDs, LLPS propensity was reduced to such an extent that we were unable to detect any droplets, even at reduced pH where LLPS is more abundant (Figs. S5–S9). For all the other variants, LLPS was more abundant at reduced pH, as expected.

Changes in LLPS propensity due to the patient mutations E83Q and A18T could likewise be rationalized by their effects on sequence complexity. Upon induction of LLPS *in vitro*, the E83Q mutation did not affect droplet formation significantly when compared to WT (Fig. 2), consistent with its minimal effect on sequence complexity. In contrast, the A18T mutation significantly reduced LLPS relative to WT (Fig. 2), similar to the effect of V16I. However, V16I had a stronger effect on LLPS compared to A18T, despite having the same sequence complexity (of 0.912), which indicates that residue-specific factors such as physicochemical properties and/or structural propensities also contribute to LLPS, in addition to sequence complexity. Nevertheless, the significant reduction of α-syn LLPS by the complexity-enhancing PD-associated mutation A18T implicates sequence complexity in the pathogenicity of α-syn.

We did not observe any changes in droplet appearance for any of the α-syn variants, on imaging at longer (than 4 hr) timepoints (Fig. S11A). This is consistent with the fact that α-syn droplets forming under our experimental conditions mature very quickly (Fig. S2). However, there was a steady increase in turbidity of all LLPS variants over time (Fig. S11B), which can be attributed to the ongoing fibril formation that is known to follow phase separation of α-syn (11; 22; 30).

Therefore, by making subtle complexity-enhancing changes to the sequence of α-syn, we were able to control its LLPS over a wider dynamic range than possible with previously studied variants (*i*.*e*., E46K and A53T (11), neither of which affects sequence complexity). We next sought to use our designed (and patient) variants to investigate the relationship between LLPS and fibril formation of α-syn.

### Relationship between LLPS and fibrillation of α-syn

LLPS of α-syn is followed by a liquid-to-solid transition that includes the formation of amyloid fibrils (fibrillation), which can be monitored by the thioflavin T (ThT)-binding assay (11; 22; 30). When we monitored the increase in ThT fluorescence over time following LLPS of each variant at neutral pH (Fig. S12), we found that the variants with greater propensity to phase separate formed fibrils faster, as indicated by their shorter half-times of fibrillation (Fig. 3). The positive correlation between rate of fibrillation and propensity to phase separate suggests that LLPS can be a key determinant of amyloid formation under conditions promoting phase separation (*i*.*e*., in presence of PEG, ‘LLPS conditions’). This correlation was robust to changes in pH (Fig. S13), further supporting the capacity of LLPS to promote fibrillation. Interestingly, despite exhibiting no measurable LLPS at any pH, the –LCD1&2 variant eventually formed fibrils in the presence of PEG, which demonstrates that LLPS is not required for fibrillation of α-syn even under conditions that promote phase separation.

**Figure 3.**
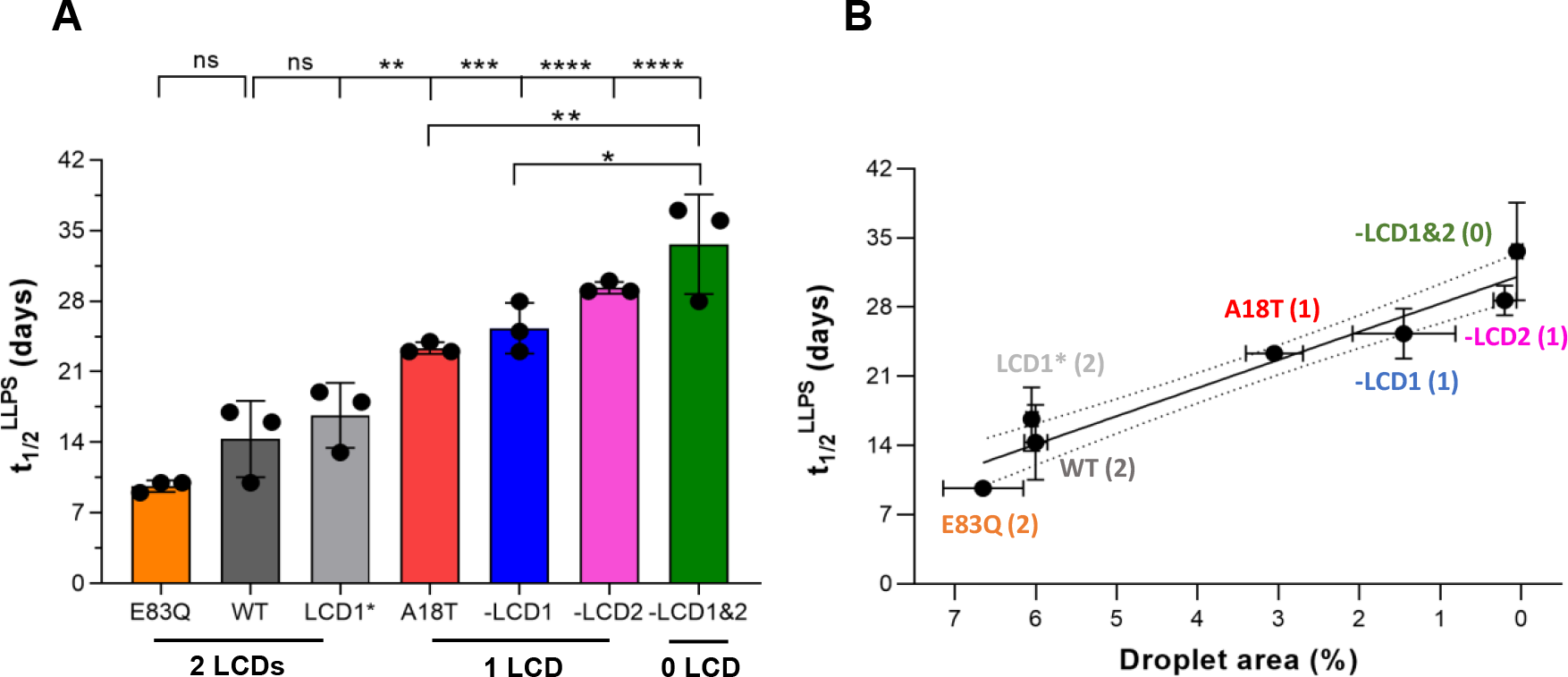
**(A)** Fibrillation half-times of different α-syn variants subjected to aggregation under LLPS conditions. **(B)** The same plotted against droplet forming propensity of the variants, showing a negative correlation between the two – variants forming more droplets forms fibrils faster (shorter half-times). Numbers in brackets represent number of LCDs. Dotted lines represent 95% confidence interval for linear regression. Data represent mean ± standard deviation (SD) for n = 3 independent experiments. Statistical significances are indicated by *: p ≤ 0.05, **: p ≤ 0.01, ***: p ≤ 0.001, ****: p ≤ 0.0001.

We next asked if mutations designed to selectively perturb LLPS also affect its fibrillation in conditions that do not promote phase separation. We therefore monitored the fibrillation kinetics of each α-syn variant in the absence of PEG, where fibrillation was induced by the conventional method of continuous shaking (‘non-LLPS conditions’). From the corresponding ThT-binding profiles (Fig. S14) and calculated half-times of fibrillation (Fig. 4A), we observed no correlation between the rate of fibrillation and either LLPS propensity (Fig. 4B) or sequence complexity (Fig. S15B). For example, whereas WT and LCD1* (with the same sequence complexity) have similar propensities to phase separate, they differ dramatically in rate of fibril formation. In contrast, whereas WT and –LCD2 have very different sequence complexities and propensities to phase separate, they form fibrils at similar rates. Therefore, by modifying sequence complexity, we could identify mutations that selectively perturb LLPS *or* fibrillation of α-syn, which might allow dissection of their individual roles *in vivo*. Interestingly, the sporadic PD variant A18T (and designed variant –LCD1) accelerated α-syn fibrillation, despite significantly *reducing* droplet formation. Also, the variant lacking both LCDs, despite being unable to phase separate, did form fibrils, albeit slower than the WT protein. These results indicate that the intrinsic determinants of α-syn LLPS and fibrillation are different – while LLPS is highly sensitive to the sequence complexity of the protein, fibrillation is not.

**Figure 4.**
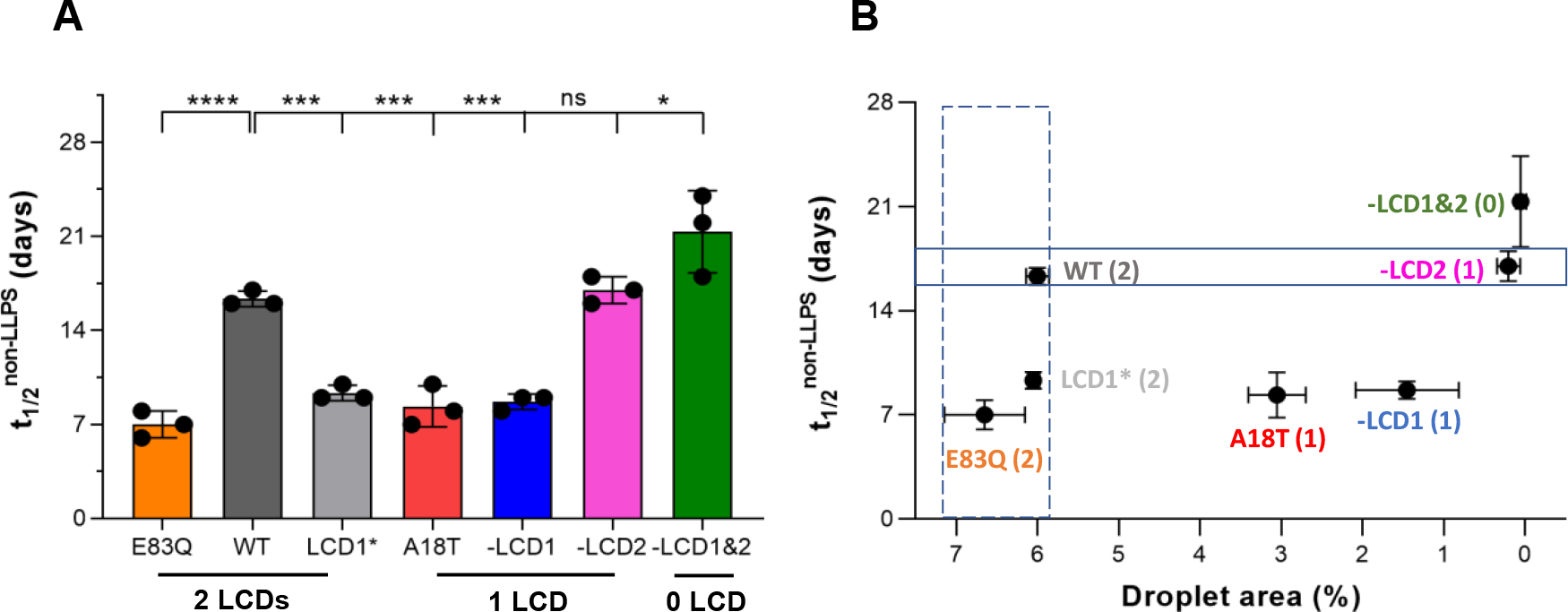
**(A)** Fibrillation half-times of different LLPS variants subjected to aggregation under non-LLPS conditions. **(B)** The same plotted against droplet forming propensity of the LLPS variants, showing a lack of correlation between the two. Numbers in brackets represent number of LCDs. The continuous box encloses variants that have similar fibrillation propensities despite drastically different phase-separating tendencies, while the box with broken lines highlights variants with different fibrillation propensities despite similar phase-separating tendencies. Data represent mean ± standard deviation (SD) for n = 3 independent experiments. Statistical significances are indicated by *: p ≤ 0.05, **: p ≤ 0.01, ***: p ≤ 0.001, ****: p ≤ 0.0001.

### LLPS-dependent *vs*. -independent fibrillation of α-syn

Recent reviews have suggested that LLPS-induced fibrillation of α-syn is an alternate mode of amyloidogenesis, which might lead to fibrillar strains that are distinct from those formed in the absence of LLPS (28; 47). We provide direct evidence for this divergence by comparing the morphology and proteolytic fingerprints of WT α-syn fibrils obtained under LLPS and non-LLPS conditions, which differ only in the presence of PEG (Fig. 5). Not only did the two types of fibrils exhibit distinct morphologies (Fig. 5A), they also produced different proteolytic fingerprints upon digestion with proteinase K (Fig. 5B). Amyloids produced following LLPS were also significantly less resistant to digestion, indicating a more solvent-exposed structure. The formation of structurally distinct amyloids in the presence *vs*. absence of LLPS would require distinct assembly mechanisms, which is consistent with the lack of correlation between the rates of fibrillation *via* these two pathways (Figs. 2 and 3). The significant molecular differences between LLPS and fibrillation further motivates efforts to dissect their individual roles in disease.

**Figure 5.**
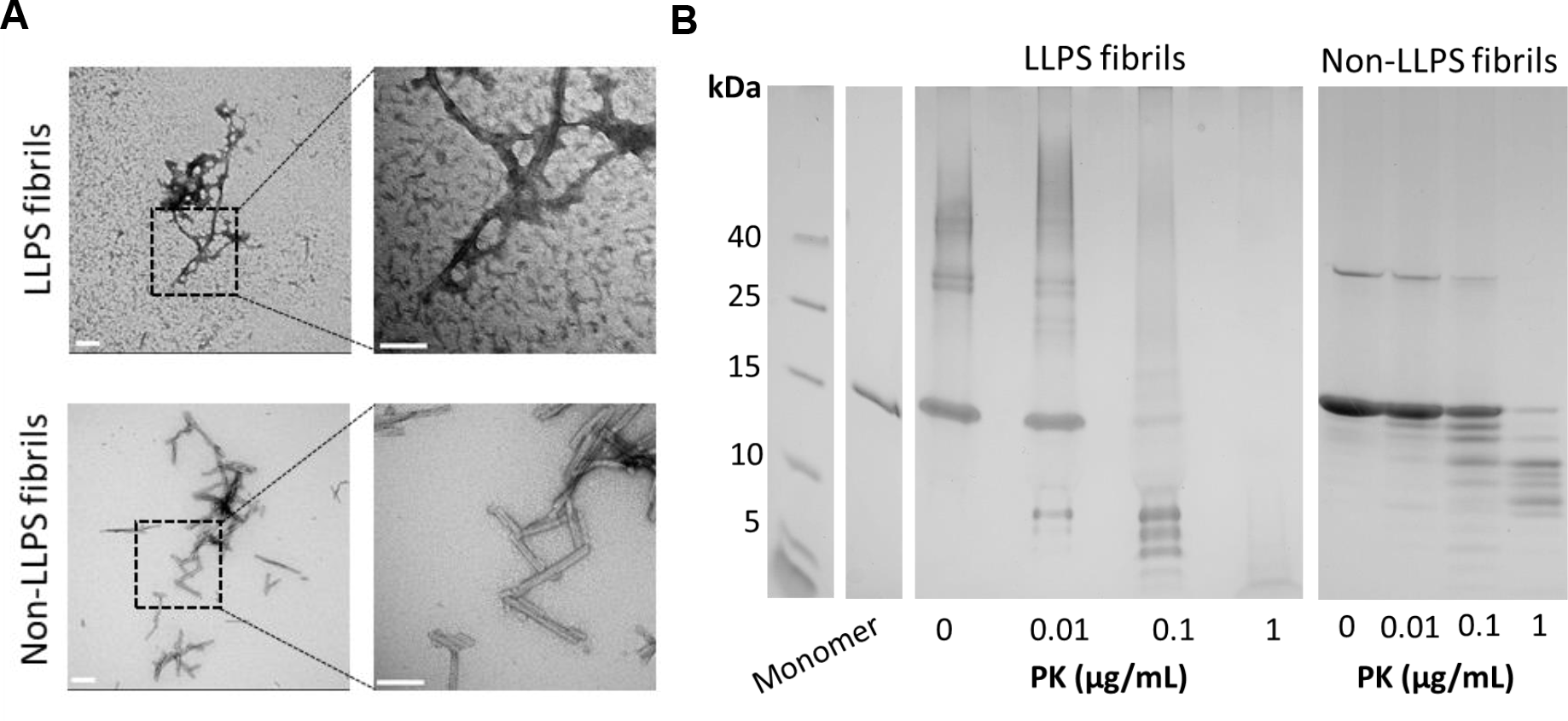
**(A)** TEM images and **(B)** proteolytic fingerprints, of WT α-syn fibrils produced under LLPS and non-LLPS conditions. Scale bar = 100 nm.

## Discussion

LLPS, which involves the metastable de-mixing of macromolecules into liquid-like droplets, has recently been implicated as a regulator of fibril formation of amyloidogenic proteins associated with disease (4). α-Syn is an amyloidogenic protein whose fibril formation is known to contribute to the pathology of PD and other synucleinopathies. Therefore, since the discovery of its propensity to undergo LLPS (11; 22), there has been significant interest in addressing the potential role of α-syn LLPS in fibrillation. A gradual liquid-to-solid transition into fibrillar aggregates has been shown to follow α-syn LLPS, and a variety of perturbations have been found to simultaneously enhance fibrillation and LLPS of α-syn – suggesting LLPS to be is an alternate nucleation mechanism in the fibrillation pathway (28). However, the intrinsic drivers of LLPS *itself* remain largely unknown, without which it is difficult to ascertain its biological significance. Moreover, the lack of strategies to selectively perturb α-syn LLPS complicates the dissection of the individual roles of LLPS and fibrillation in disease.

In this paper, using a combination of carefully designed mutations that target the two LCDs of α-syn, we demonstrate that low sequence complexity is one key intrinsic driver of α-syn LLPS. To test the role of sequence complexity in α-syn LLPS, we introduced chemically subtle, complexity-enhancing point mutations in the two LCDs of α-syn (to make the protein sequence progressively more complex without altering its physicochemical or structural properties), and studied the ability of the resulting variants to phase separate relative to the WT. We found that α-syn’s propensity to phase separate decreases with increasing sequence complexity, which was also reflected in relevant patient mutations (Fig. 2). Note that the dramatic reduction in α-syn LLPS by the extremely subtle, complexity-enhancing V16I mutation (in variant –LCD1), and its reversal by another adjacent V15I mutation (in variant LCD1*), underscores the exquisite sensitivity of α-syn phase separation to sequence complexity. Although the role of LCDs in protein LLPS has typically been attributed to their unique amino acid compositions (4), our results demonstrate that the low sequence complexity of the LCDs can *itself* be a driver of LLPS.

Furthermore, leveraging the differences in LLPS propensities among different α-syn variants, we demonstrate that fibrillation of α-syn does not necessarily correlate with its propensity to phase separate (Fig. 4B). In doing so, we identify mutations that selectively perturb α-syn LLPS (–LCD2) *or* fibrillation (LCD1*), which should allow dissection of their individual roles *in vivo*. It is interesting to note that previously studied fibrillation-promoting patient mutations of α-syn (*i*.*e*., E46K and A53T), were both found to increase LLPS (11), suggesting a pathological role of α-syn LLPS in PD. However, in this work, we identified two patient variants – E83Q and A18T, which do not increase LLPS despite accelerating fibrillation; in fact, A18T *reduces* LLPS. This indicates that the role of LLPS in pathological fibrillation of α-syn, and hence in PD, might be more complex than previously appreciated.

Overall, our results establish that low sequence complexity is a key driver of α-syn phase separation but not necessarily fibrillation. We speculate that low sequence complexity contributes to LLPS by enabling a variety of distinct intermolecular contacts with similar energy, promoting dynamic self-association and preventing the rapid adoption of a unique and stable conformational state. Similar phenomena have been implicated in molten globule proteins that likewise involve protein condensation without the formation of a unique structure (48). Conversely, increasing sequence complexity might reduce the number of intermolecular contacts that have similar energy, preventing the coexistence of many conformational states, which might be required for dynamic self-assembly. Regardless of the molecular mechanism, our results highlight actionable design principles for controlling LLPS using sequence complexity, which can also be applied to decouple LLPS and fibrillation. We therefore believe that the variants we generated by these principles will enable future studies to distinguish the cellular/*in vivo* effects of α-syn LLPS from those of its amyloid formation, and can also form the basis for disentangling LLPS and amyloid formation of other phase-separating amyloidogenic proteins similar to α-syn.

## Materials and Methods

### Computational tools used in the study

To design mutations in the low-complexity domains (LCDs) of α-syn that would not alter physicochemical or structural properties of the protein, we selected substituent amino acids that have the same charge as the ones being replaced (neutral), and ensured that their hydrophobicity and α-helix preferences were as close as possible to the ones being replaced. The overall hydrophobicity and charge of each resulting variant were calculated using the GRAVY (*www.gravy-calculator.de*) and Prot pi (*www.protpi.ch/Calculator/ProteinTool*) online calculators respectively. The helix-forming propensity was obtained from the α-helix calculator by Deleage & Rouge (49) in ProtScale (*web*.*expasy*.*org/protscale*), and disorder was calculated using IUPred2A (50) (*iupred2a*.*elte*.*hu*). After ensuring that the introduced mutations did not significantly alter any of these properties of the protein, their effects on sequence complexity were checked using the SMART prediction tool (42) (*smart*.*embl-heidelberg*.*de*). Disease mutations were also similarly analysed.

### Design strategy of LLPS variants using SMART

SMART (Simple Modular Architecture Research Tool) is a freely available software that engages the well-known SEG algorithm(43-45) to predict LCDs in protein sequences. The SEG algorithm classifies protein segments as LCDs based solely on their compositional complexity. Compositional complexity of a protein segment is determined by the count of unique amino acids in that specific segment, irrespective of sequence patterns or repetition (45). Mathematically, compositional complexity of a subsequence of length *L*, is defined by 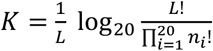, where *n*_*i*_ is the number of occurrences of each of the 20 different amino acids in the given segment, with 0≤*n*_*i*_≤*L* and 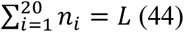. As the number of distinct amino acids in a given segment of length *L* increases, one or more *n*_*i*_ values decrease, resulting in a smaller denominator within the logarithmic term in *K*. Consequently, the logarithmic term, and hence the compositional complexity of the segment, increases with an increasing number of distinct amino acids in that segment. Importantly, the compositional complexity of a given segment remains the same for the same set of *n*_*i*_values (representing the count of different amino acids) regardless of the specific identities of the amino acids (*i*).

To test the role of sequence complexity in α-syn LLPS, we designed subtle missense mutations in the two LCDs of the protein (LCD1 and LCD2), such that they increase sequence complexity of the protein while minimally perturbing its structural and physicochemical properties. The LCD1 region of WT α-syn (residues 10-23) is composed of six different amino acids – **four** Ks, **four** As, **two** Es, **two** Vs, **one** G, and **one** T. Accordingly, *K* for this region is 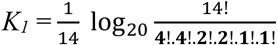 Replacing one of the six different amino acids in LCD1 with an amino acid that doesn’t exist in this segment is expected to increase compositional complexity of the segment by increasing the number of distinct constituent amino acids (and consequently lowering the value of the denominator within the logarithmic term in *K*_*1*_). Accordingly, we introduced a single V-to-I (valine-to-isoleucine) mutation in this region (V16I); and found it sufficient to increase sequence complexity of this region above the threshold for identification as a LCD by SMART (Fig. 1, variant –LCD1). We chose a V-to-I mutation in particular as it introduces an extremely subtle change in the protein (adds a single methyl group on a single hydrophobic residue in a 140-residue protein), which should minimally perturb the structural and physicochemical properties of the protein. This was confirmed from computational analyses, which predicted minimal changes to the protein’s charge and hydrophobicity (Table S1), as well as its propensity for disordered and helical structures (Figs. S1A and C). Note that changing one of the two valines in LCD1 to isoleucine (in case of V16I) increases *K*_*1*_ by lowering the denominator within the logarithm; however, changing both of the two valines to isoleucines in this region would restore the original value of the denominator and consequently *K*_*1*_ (same set of n_i_ values as for the WT protein). Therefore, to test if restoring the original sequence complexity can rescue LLPS behaviour of the WT protein, we designed another variant in which the increased local complexity of the LCD1 region by the V16I mutation was reversed by introducing another V-to-I mutation at position 15 (Fig. 1, variant LCD1*). As a result, this variant had the same sequence complexity as the WT protein, but a slightly different amino acid sequence (with two Is instead of two Vs at positions 15 and 16 within the LCD1 region).

The LCD2 region of WT α-syn (residues 63-78) has five different amino acids: six Vs, three As, three Gs, three Ts, and one N. Accordingly, *K* for this region is: 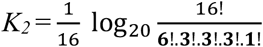 Following the same principle as that used for designing the –LCD1 variant, we first introduced a single V-to-I mutation at position 70, to increase complexity of this region (by lowering the value of the denominator within the logarithmic term in *K*_*2*_). However, a single substitution was not sufficient to increase complexity of this relatively longer LCD above the threshold for detection as a LCD by SMART. Therefore, we paired another G-to-P mutation at position 68 to lower *K*_*2*_ further. The combined changes (G68P and V70I) were sufficient to increase complexity of the LCD2 region above the threshold for identification as a LCD by SMART (Fig. 1, variant –LCD2). Then we combined the –LCD1 and –LCD2 substitutions to make a variant with no predicted LCDs (Fig. 1, variant –LCD1&2).

### Quantification of sequence complexity of LLPS variants

For the purpose of our study, we defined sequence complexity of α-syn (S) as the sum of the local compositional complexities of the two segments corresponding to its two LCDs. Therefore S = *K*_*1*_ *+ K*_*2*_. Note that for the E83Q variant of α-syn used in our study, the second LCD spans residue 63-83 instead of 63-78 as for the WT (see Fig. 1). Hence, for the sake of maintaining uniformity in our quantification parameter, we used the local compositional complexity of this stretch as our *K*_*2*_ in the expression of S.

Therefore S = *K*_*1*_ *+ K*_*2*_, where *K*_*1*_ and *K*_*2*_ are the local compositional complexities of segments 10-23 and 63-83 in each of the α-syn variants used in our study.

Accordingly, the calculations of sequence complexities for the variants used in our study, have been tabulated below:

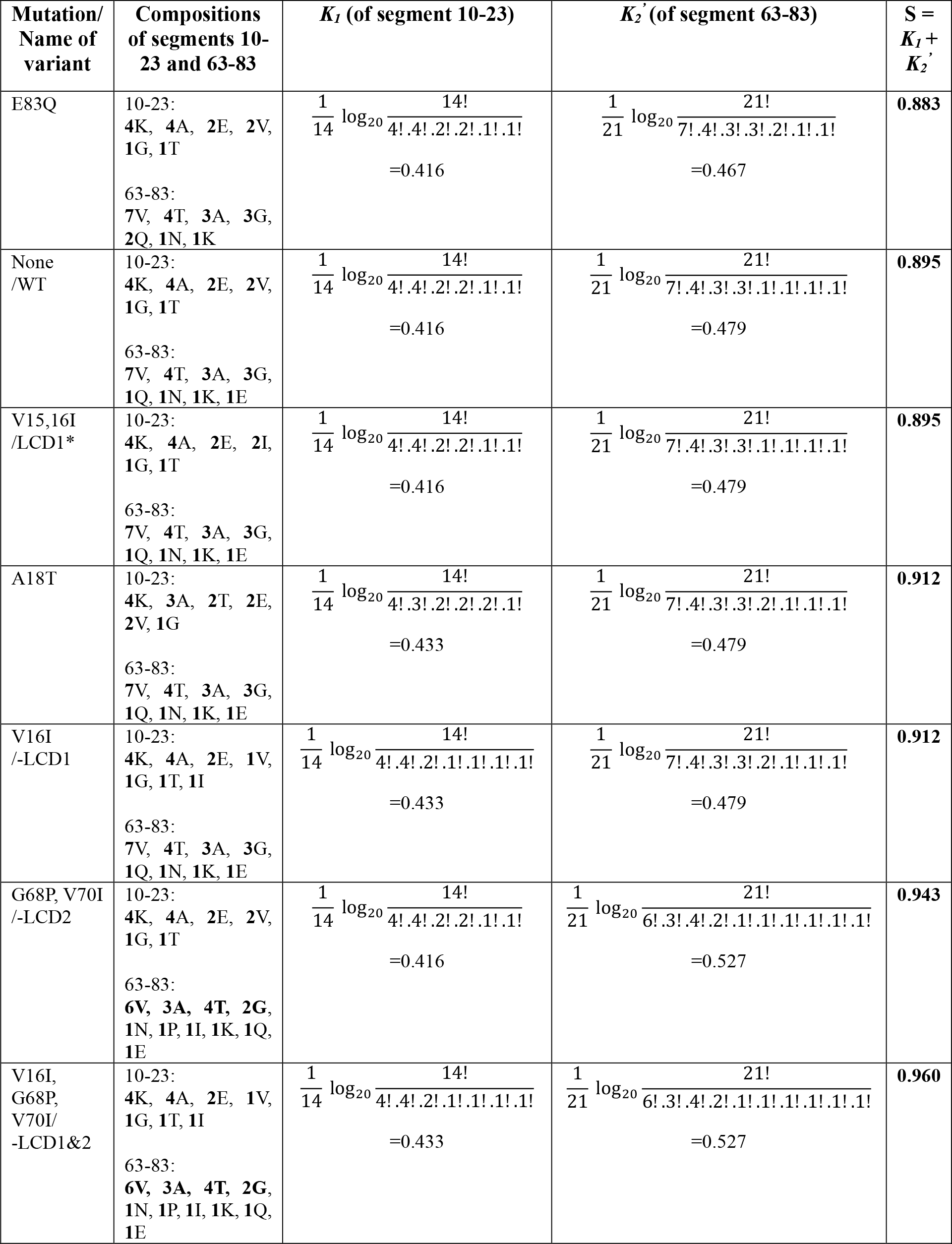

### Site-directed mutagenesis (SDM)

The recombinant bacterial plasmid pET-28a, containing a kanamycin resistant gene and the gene for α-syn, was used as a template for SDM to produce all the mutants apart from –LCD1&2, which was obtained from Twist Bioscience (USA). For every other mutant, mutagenic primers were designed (see below) and the mutagenesis was done using either the Phusion High-Fidelity PCR Kit (ThermoFisher Scientific, USA) or the QuikChange Lightning Site-Directed Mutagenesis Kit (Agilent Technologies, USA), following respective manufacturers’ protocols. The Simply Amp Thermal Cycler (ThermoFisher Scientific, USA) was used to carry out all PCR. When using the Phusion kit, DpnI digestion of the PCR product was done using 1μL of DpnI added to 50μL reaction mixture and incubating for 1 hour at 37 °C. This was followed by PCR clean-up using the GeneJet PCR Purification Kit (ThermoFisher Scientific, USA), following manufacturer’s protocol. The resulting DNA was then transformed into competent DH5-alpha cells, which were plated on kanamycin selective LB-agar plates and incubated overnight for growth of colonies containing the desired mutation. For PCR using the Agilent kit, DpnI digestion and subsequent transformation into XL 10-Gold Ultracompetent Cells was done using reagents provided in the kit, following manufacturer’s protocol. Single colonies resulting from successful PCR and transformations were grown overnight in 5 mL cultures, and DNA from them were extracted using the GeneJET Plasmid Miniprep kit following manufacturer’s protocol. The resulting mutations were confirmed by Sanger sequencing of the extracted DNA samples (through Eton Bioscience).

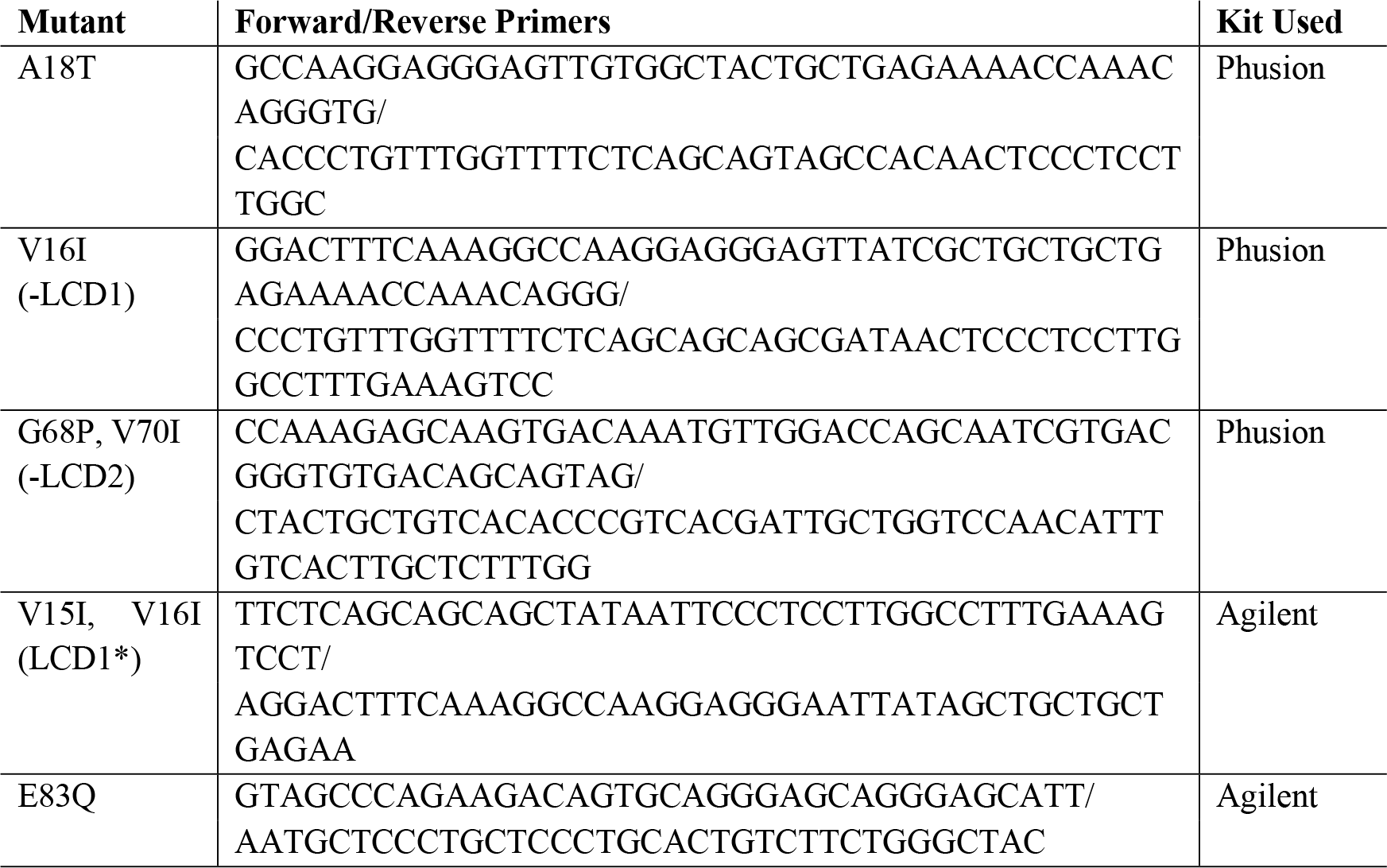

The primers used for the SDM reactions are as follows:

### Expression and purification of α-syn variants

Recombinant WT α-syn and its mutants were individually expressed in the *Escherichia coli* BL21 (DE3) cells transformed with the respective plasmids. Expression was induced using 1 mM of isopropyl β-D-1-thiogalactopyranoside (IPTG) at an OD of 0.5–0.6. The cultures were incubated at 37 °C with shaking at 225 rpm for 4 hours after addition of IPTG. Cells were then harvested by centrifugation, and stored at -80 °C until purification of α-syn by a previously reported non-chromatographic method (24; 51). Briefly, the cell pellets were resuspended in Tris buffer (50 mM Tris, 10 mM EDTA and 150 mM NaCl, pH 8) and boiled for 10 minutes, after adding 100 mM phenylmethylsulfonyl fluoride (PMSF). Lysed cells were then centrifuged and the supernatant was removed into a fresh tube. Streptomycin sulfate (136 μL of 10% solution per mL supernatant) and acetic acid (glacial, 228 μL per mL supernatant) were added to it, followed by additional centrifugation at 12000 g for 5 minutes at 4°C. The supernatant was removed again and then precipitated with solid ammonium sulfate (up to 50% saturation, as calculated using the online Ammonium Sulfate Calculator from EnCor Biotechnology Inc.) at 4°C. The precipitated protein was collected by centrifugation for 30 minutes at 12000 g, and the pellet was washed once with 50% ammonium sulfate solution. The washed pellet was re-suspended in minimum volume of 100 mM ammonium acetate (to form a cloudy solution) and precipitated by adding an equal volume of 100% ethanol at room temperature. Ethanol precipitation was repeated once more, followed by freezing of the obtained pellet, and subsequent lyophilization using a FreeZone lyophilizer by Labconco (USA).

### Preparation of Low Molecular Weight (LMW) forms of α-syn variants

LMW forms of all variants were prepared using a previously published protocol (11), with slight modifications. Lyophilized protein was dissolved in 20 mM sodium phosphate buffer (NaP) at pH 7, at ∼20 mg/mL. If required, the protein was solubilized by the addition of a few drops of 0.2 N NaOH and the final pH was adjusted to 7 using 2 M HCl. The protein solution was then centrifuged at 15000 rpm for 30 minutes at 4 °C to remove insoluble aggregates. Thereafter, the supernatant was passed through a pre-washed 100 kDa cut-off filter (Amicon Ultra-0.5 Centrifugal Filter Unit) to remove any high-order aggregates. The flow-through *i*.*e*., the LMW fraction constituting majorly of monomeric α-syn was collected, and the concentration was estimated by measuring the absorbance at 280 nm. The molar extinction coefficient (ε) used for calculating concentrations of all variants was 5960 M^-1^cm^-1^ (obtained using Prot pi).

### Protein labelling with NHS-Rhodamine

For the purpose of fluorescence microscopy, all samples were doped with 1% of the respective rhodamine-labelled proteins. To obtain the labelled proteins, purified variants of α-syn were labelled with NHS-rhodamine dye (ThermoFisher Scientific, USA) following the manufacturer’s protocol. Briefly, 10X molar excess of dye (dissolved in DMSO) was added to the LMW protein, and the reaction was allowed to proceed at room temperature for 2 hours in the dark. Excess unreacted dye was removed by dialysis against NaP (pH 7) at 4° C for 48 hours, with 3-4 changes of buffer.

### *In vitro* liquid-liquid phase separation (LLPS)

To induce LLPS, WT α-syn and its variants, at a concentration of 100 μM, were suspended in NaP in presence of 150 mM NaCl (PBS), 1 mM sodium azide and 20% PEG 8000, at pH 7, 6 and 5. Reaction mixtures prepared under these conditions (LLPS conditions) were added to the wells of half-area 96-well plates in triplicates, and incubated in a moist chamber at 37 °C. For droplet imaging and quantification, samples were prepared with 1% of labelled proteins, and incubated for up to 4 hours. For monitoring fibril formation by the thioflavin T (ThT)-binding assay (*see section* ***S1*.*9***), samples without labelled proteins were incubated in presence of 25 μM ThT under the same conditions.

### Fluorescence microscopy and image quantification

Droplet formation by LLPS variants of α-syn was visualized under a ‘EVOS FL Auto 2’ fluorescence microscope (ThermoFisher Scientific, USA), in the fluorescence mode, using the EVOS™ Light Cube, RFP 2.0 (excitation: 542/20, emission: 593/40). Appropriate buffer controls for each experiment were kept for baseline fluorescence settings, and all the images were obtained at a resolution of 2080 x 1552 pixels at 32-bit depth. Samples were either viewed through the LWD EVOS™ 10X Objective (fluorite, NA = 0.3, WD = 7.13) or the LWD EVOS™ 60X Objective (fluorite, NA = 0.75, WD = 2.2 mm), for capturing low and high magnification images respectively.

Quantification of droplet formation was done from the low magnification (10X) images of 4-hour samples, using Fiji (ImageJ). For every image, at first the scale was set for particle size measurements using the scale bar from the microscope. Then the image was converted to an 8-bit image, followed by thresholding to include only droplets in the particle size measurements. The sizes (areas) of all highlighted droplets in the field of view were then measured, using the ‘Analyse particles’ module, setting the lower limit of size at 3.14 μm^2^ (radius = 1 μm) and keeping circularity within 0-1. Individual areas thus obtained were used to estimate diameters of the droplets and plot the corresponding size distributions. Fractional area covered by droplets and number of droplets were also obtained from the analyses, which were multiplied to obtain the percentage of total area covered by droplets in each of the samples.

### Confocal microscopy and Fluorescence Recovery After Photobleaching (FRAP)

FRAP studies were performed using a laser scanning confocal microscope (Zeiss LSM 710, inverted) equipped with a Plan-Apochromat 63X/1.4 NA oil immersion objective (WD = 0.19 mm). After 1 hour of incubation of all α-syn variants under LLPS conditions, droplet formation was first confirmed by fluorescence microscopy (EVOS FL Auto 2), and then the wells containing droplets were subjected to FRAP experiments in the confocal microscope. Intensity was recorded from three different region of interests (ROIs) of constant radii – actual bleaching region (ROI-1), reference region on same/neighboring droplet (ROI-2) at a different location to correct for passive bleaching during laser exposure, and a region outside the droplet in the dark (ROI-3) to correct for background fluorescence intensity. The selected ROI-1 was then bleached with 50% laser power, using a 561 nm DPSS 561-10 laser and emission intensities from all ROIs were recorded for 100s (when fluorescence emission from ROI-1 reached a plateau for the WT variant). All measurements were performed at room temperature.

FRAP data analyses were performed according to previous studies (11; 52). First, all fluorescence recovery curves were constructed from the total intensity values in the ROI for each frame corrected for the background and passive bleaching. To calculate corrected and normalized fluorescence intensity, I (n), the following equation (Eq. 1) was used:

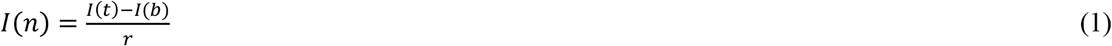

Where, I (t) = fluorescence intensity at time t, I (b) = background fluorescence intensity and rate of photo-bleaching 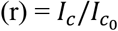

[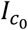 = fluorescence intensity of the ROI-2 before photo-bleaching; *I*_*c*_ = fluorescence intensity of the ROI-2 after photo-bleaching]

The normalized and background corrected fluorescence recovery curves thereby obtained were fitted using the following single exponential recovery function (Eq. 2) in the OriginPro8.5 software (OriginLab, USA):

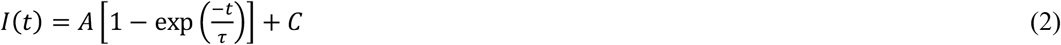

Where, τ is the fluorescence recovery time constant, ‘A’ corresponds to the mobile fraction of the fluorescent probe and ‘C’ is the y-intercept of the recovery curve. Note that mobile fraction (M.F) is depicted by: 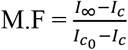

### Thioflavin T (ThT)-binding assay

ThT-binding assay was used for monitoring the fibrillation of all variants of α-syn in the presence and absence of LLPS. For fibrillation under LLPS conditions, all variants at a concentration of 100 μM, were suspended in NaP in presence of 150 mM NaCl, 1 mM sodium azide, 25 μM ThT and 20% PEG 8000, at pH 7, 6 and 5. Samples were prepared and loaded in triplicates into half-area 96-well plates, and incubated under quiescent conditions in a moist chamber at 37 °C. For fibrillation under non-LLPS conditions, all variants at a concentration of 100 μM, were suspended in NaP in presence of 150 mM NaCl and 25 μM ThT, at pH 7. Samples were prepared and loaded in triplicates into half-area 96-well plates, and shaken continuously at 800 rpm at 37 °C in a ThermoMixer C (Eppendorf, Germany). At regular time intervals, the plates were read using a Synergy H1 microplate reader from BioTek (USA), to monitor the changes in ThT fluorescence over time, usually for up to a month. Since all replicates of the -LCD1&2 variant under LLPS conditions did not show saturation of ThT signal within a month, they were monitored for an additional two weeks until saturation. The steady state fluorescence of ThT was measured using excitation at 440 nm and emission at 485 nm. The resulting ThT-binding profiles were then fit into the Boltzmann equation (Eq. 3) as follows:

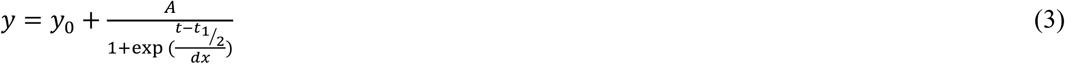

Where *y*_*0*_ is the signal baseline at the beginning, *A* is the total increase in fluorescence signal, (*1/dx*) is the growth rate constant, and *t*_1/2_ is the mid-point of the transition *i*.*e*., half-time of fibrillation.

Curve fitting was done in the OriginPro8.5 software (OriginLab, USA), and corresponding half-times were calculated for further analyses.

### Preparation of pre-formed fibrils (PFFs) of α-syn

To prepare WT α-syn fibrils under LLPS conditions, the un-labelled LMW protein was incubated at a concentration of 100 μM in NaP at pH 7, in presence of 150 mM NaCl, 1 mM sodium azide and 20% PEG 8000, for a month. To prepare α-syn fibrils under non-LLPS conditions, the un-labelled LMW variants were incubated at a concentration of 100 μM in NaP at pH 7, in presence of 150 mM NaCl, under constant agitation at 800 rpm in a ThermoMixer C (Eppendorf, Germany), for a month. Fibril formation in the resulting turbid solutions was checked by ThT-binding fluorescence, and samples were centrifuged at 15000 rpm for 30 minutes to pellet down fibrils. Clear supernatants from the top were carefully removed and their volumes determined. For every non-LLPS fibril sample, the protein concentration of the supernatant was measured by recording absorbance at 280 nm (OD_280_); and quantity of fibrils formed was determined (in micrograms) by subtracting the amount of protein left in the supernatant (volume times concentration) from the initial amount of protein. The pellets were then resuspended in required volumes of NaP (pH 7) to obtain a final concentration of 1 mg/mL for each. For the WT fibril sample produced under LLPS conditions, determining protein concentration by OD_280_ measurement was not feasible due to absorption of PEG at 280 nm. Therefore, fibril pellet was resuspended in a volume of NaP (pH 7) that was equal to the supernatant volume, so as to obtain ≤ 1.4 mg/mL (100 μM) of LLPS fibrils. All samples were aliquoted and stored at -80 °C until further use.

### Transmission Electron Microscopy (TEM)

5 μl of PFF solution (∼1 mg/mL) was spotted on a freshly glow-discharged 300-mesh carbon coated formvar grid (Electron Microscopy Sciences, USA) and incubated for 3 minutes. Excess solution was soaked off using a filter paper, and the grid was then washed twice Milli-Q water. 5 μl of 1% (w/v) uranyl acetate solution was added onto the grid and allowed to sit for 2 minutes before soaking off excess with a filter paper. All grids were air-dried, and imaged using a Tecnai Biotwin Spirit Transmission Electron Microscope (TEM), operated at 80 kV. Images were captured using an AMT 1kx1k camera at 43000X and 135000X magnifications, using the TIA imaging system provided with the instrument.

### Preparing LLPS fibrils for proteolytic fingerprinting

Proteolytic fingerprinting of fibrils involves subjecting fibril samples to limited proteolysis by Proteinase K (or PK) followed by running a SDS-PAGE with digested samples to identify the characteristic band patterns (position and relative intensity) obtained on digestion. As presence of PEG in LLPS fibril samples interferes with running of the protein gel and creates smeared bands (data not shown), these particular fibril samples were subjected to an additional wash step prior to being used for proteolytic fingerprinting experiments. After isolating the LLPS fibrils as pellets using high-speed centrifugation, the obtained pellet was resuspended in 500 μL of NaP buffer, and passed through a pre-washed 100 kDa cut-off spin column (Amicon Ultra-0.5 Centrifugal Filter Unit). The residual volume of sample retained in the spin column, containing fibrillar aggregates (>100 kDa), was then washed twice more with 500 μL of buffer. Finally, the fibrillar aggregates retained in the spin column, free of most of the PEG contamination after three wash steps, were recovered by inverting the spin column and centrifuging it once more. The sample thus obtained was then subjected to proteolytic fingerprinting experiments (see below).

### Limited proteolysis using Proteinase K (PK)

Limited proteolysis of protein samples with PK generates distinct peptide fragments depending on which epitopes of the protein are accessible to the enzyme. Suitable amounts of WT α-syn PFFs formed under LLPS and non-LLPS conditions were mixed with 0, 0.01, 0.1 and 1 μg/ml of PK (20 mg/mL) in NaP to a final volume of 12 μl, and incubated at 37°C for 10 minutes. PK digestion was stopped with 1 mM PMSF. Reaction samples were then boiled with equal volumes of Novex™ Tricine SDS Sample Buffer (2X) at 85 °C for 2 minutes, and then centrifuged at high speed for 5 minutes. Samples were then run and resolved on Novex™ 16% Tricine gels (1.0 mm, Mini Protein Gels) supplied by Invitrogen, to obtain respective proteolytic fingerprints. Fibrillar samples of all mutants of α-syn obtained under non-LLPS conditions were also digested with 1 μg/ml of PK for 10 minutes, and processed similarly, for identifying the relative differences in their proteolytic fingerprints.

### Data plotting and statistical data analyses

All experimental data were plotted and fit using OriginPro 8.5 or GraphPad Prism 9 software. All statistical data analyses were performed using one-way ANOVA (Tukey’s post hoc) in the GraphPad Prism 9 software, where ns: nonsignificant, *: *p* ≤ 0.05, **: *p* ≤ 0.01, ***: *p* ≤ 0.001, and ****: *p* ≤ 0.0001.

## Supporting Information

Supplementary figures, table and text are included in the supporting information. Supplementary figures: disorder and helix score analyses of all α-syn variants, droplet formation and maturation over time for WT α-syn, turbidity values of all α-syn variants at pH 7 after 4 hr of incubation, LLPS propensity of the V70I variant, size distribution and droplet area of all α-syn variants at different pH, low and high magnification fluorescence microscopic images of all α-syn variants at different pH, fluorescence recovery after photobleaching (FRAP) of droplets, high magnification images and turbidity values at different time points for all α-syn variants, representative ThT-binding profiles of all α-syn variants under LLPS and non-LLPS conditions, fibrillation half-times *vs*. droplet area and sequence complexity of all α-syn variants under LLPS and non-LLPS conditions, proteolytic fingerprinting of LLPS and non-LLPS fibrils of WT α-syn. Supplementary table: calculated hydrophobicity scores and charges of all α-syn variants. Supplementary text: results of FRAP experiments on droplets of α-syn variants.

## Author Contributions

A.M. conceived the project. A.M. and R.W.N. designed the experiments and data analyses, which were performed by A.M. The manuscript was written and approved in its final form by both authors.

## Conflicts of interest

There are no conflicts to declare.

## Acknowledgements

This work was supported by grants from the NIH (R00-NS116679) and the Welch Foundation (F-2116-20220331). Transmission electron microscopy and confocal microscopy were performed at the Center for Biomedical Research Support (CBRS) Microscopy and Imaging Facility at UT Austin (RRID:SCR_021756).

## Notes

### Competing Interest Statement

The authors have declared no competing interest.

### Summary of Updates

'Introduction' and 'Materials and Methods' sections updated to clarify sequence complexity and design strategy of variants used. 'Results and Discussion' section modified to include additional supporting data. Figures 1, 2, 3, and 4 revised, and Figure 5 added in the manuscript, according to changes made.

## References

1. Hyman AA, Weber CA, Jülicher F. Liquid-liquid phase separation in biology. Annu Rev Cell Dev Biol. 2014;30:39–58.

2. Dobson CM. The amyloid phenomenon and its links with human disease. Cold Spring Harb Perspect Biol. 2017;9:a023648.

3. Eisenberg D, Jucker M. The amyloid state of proteins in human diseases. Cell. 2012;148:1188–1203.

4. Zbinden A, Pérez-Berlanga M, De Rossi P, Polymenidou M. Phase separation and neurodegenerative diseases: a disturbance in the force. Dev Cell. 2020;55:45–68.

5. Babinchak WM, Surewicz WK. Liquid–liquid phase separation and its mechanistic role in pathological protein aggregation. J Mol Biol. 2020;432:1910–1925.

6. Lin Y, Protter DS, Rosen MK, Parker R. Formation and maturation of phase-separated liquid droplets by RNA-binding proteins. Mol Cell. 2015;60:208–219.

7. Molliex A, Temirov J, Lee J, Coughlin M, Kanagaraj AP, Kim HJ, Mittag T, Taylor JP. Phase separation by low complexity domains promotes stress granule assembly and drives pathological fibrillization. Cell. 2015;163:123–133.

8. Murakami T, Qamar S, Lin JQ, Schierle GSK, Rees E, Miyashita A, Costa AR, Dodd RB, Chan FT, Michel CH. ALS/FTD mutation-induced phase transition of FUS liquid droplets and reversible hydrogels into irreversible hydrogels impairs RNP granule function. Neuron. 2015;88:678–690.

9. Patel A, Lee HO, Jawerth L, Maharana S, Jahnel M, Hein MY, Stoynov S, Mahamid J, Saha S, Franzmann TM. A liquid-to-solid phase transition of the ALS protein FUS accelerated by disease mutation. Cell. 2015;162:1066–1077.

10. Wegmann S, Eftekharzadeh B, Tepper K, Zoltowska KM, Bennett RE, Dujardin S, Laskowski PR, MacKenzie D, Kamath T, Commins C. Tau protein liquid–liquid phase separation can initiate tau aggregation. EMBO J. 2018;37:e98049.

11. Ray S, Singh N, Kumar R, Patel K, Pandey S, Datta D, Mahato J, Panigrahi R, Navalkar A, Mehra S, Gadhe L, Chatterjee D, Sawner AS, Maiti S, Bhatia S, Gerez JA, Chowdhury A, Kumar A, Padinhateeri R, Riek R, Krishnamoorthy G, Maji SK. α-Synuclein aggregation nucleates through liquid–liquid phase separation. Nat Chem. 2020;12:705–716.

12. Johnson BS, Snead D, Lee JJ, McCaffery JM, Shorter J, Gitler AD. TDP-43 is intrinsically aggregation-prone, and amyotrophic lateral sclerosis-linked mutations accelerate aggregation and increase toxicity. J Biol Chem. 2009;284:20329–20339.

13. Conicella AE, Zerze GH, Mittal J, Fawzi NL. ALS mutations disrupt phase separation mediated by α-helical structure in the TDP-43 low-complexity C-terminal domain. Structure. 2016;24:1537–1549.

14. Zhou X, Sumrow L, Tashiro K, Sutherland L, Liu D, Qin T, Kato M, Liszczak G, McKnight SL. Mutations linked to neurological disease enhance self-association of low-complexity protein sequences. Science. 2022;377:eabn5582.

15. Zhao M, Kim JR, van Bruggen R, Park J. RNA-binding proteins in amyotrophic lateral sclerosis. Mol Cell. 2018;41:818.

16. Grundke-Iqbal I, Iqbal K, Quinlan M, Tung Y-C, Zaidi MS, Wisniewski HM. Microtubule-associated protein tau. A component of Alzheimer paired helical filaments. J Biol Chem. 1986;261:6084–6089.

17. Spillantini MG, Schmidt ML, Lee VM-Y, Trojanowski JQ, Jakes R, Goedert M. α-Synuclein in Lewy bodies. Nature. 1997;388:839–840.

18. Lee JC, Langen R, Hummel PA, Gray HB, Winkler JR. α-Synuclein structures from fluorescence energy-transfer kinetics: Implications for the role of the protein in Parkinson’s disease. Proc Natl Acad Sci USA. 2004;101:16466–16471.

19. Huang S, Mo X, Wang J, Ye X, Yu H, Liu Y. α-Synuclein phase separation and amyloid aggregation are modulated by C-terminal truncations. FEBS Lett. 2022;596:1388–1400.

20. Nelson SL, Li Y, Chen Y, Deshmukh L. Avidity-Based Method for the Efficient Generation of Monoubiquitinated Recombinant Proteins. J Am Chem Soc. 2023;145:7748–7752.

21. Ray S, Mason TO, Boyens-Thiele L, Farzadfard A, Larsen JA, Norrild RK, Jahnke N, Buell AK. Mass photometric detection and quantification of nanoscale α-synuclein phase separation. Nat Chem. 2023:1–11.

22. Hardenberg MC, Sinnige T, Casford S, Dada ST, Poudel C, Robinson EA, Fuxreiter M, Kaminksi CF, Kaminski Schierle GS, Nollen EA, Dobson CM, Vendruscolo M. Observation of an αsynuclein liquid droplet state and its maturation into Lewy body-like assemblies. J Mol Cell Biol. 2021;13:282–294.

23. Goedert M. Alpha-synuclein and neurodegenerative diseases. Nat Rev Neurosci. 2001;2:492–501.

24. Mahapatra A, Mandal N, Chattopadhyay K. Cholesterol in Synaptic Vesicle Membranes Regulates the Vesicle-Binding, Function, and Aggregation of α-Synuclein. J Phys Chem B. 2021;125:11099–11111.

25. Fusco G, Sanz-Hernandez M, De Simone A. Order and disorder in the physiological membrane binding of α-synuclein. Curr Opin Struct Biol. 2018;48:49–57.

26. Fink AL. The aggregation and fibrillation of α-synuclein. Acc Chem Res. 2006;39:628–634.

27. Goldman SM. Environmental toxins and Parkinson’s disease. Annu Rev Pharmacol Toxicol. 2014;54:141–164.

28. Mukherjee S, Sakunthala A, Gadhe L, Poudyal M, Sawner AS, Kadu P, Maji SK. Liquid-liquid phase separation of α-synuclein: a new mechanistic insight for α-synuclein aggregation associated with Parkinson’s disease pathogenesis. J Mol Biol. 2023;435:167713.

29. Kumar S, Jangir DK, Kumar R, Kumari M, Bhavesh NS, Maiti TK. Role of sporadic Parkinson disease associated mutations A18T and A29S in enhanced α-Synuclein fibrillation and cytotoxicity. ACS Chem Neurosci. 2018;9:230–240.

30. Sawner AS, Ray S, Yadav P, Mukherjee S, Panigrahi R, Poudyal M, Patel K, Ghosh D, Kummerant E, Kumar A. Modulating α-Synuclein Liquid–Liquid Phase Separation. Biochemistry. 2021;60:3676–3696.

31. Xu B, Huang S, Liu Y, Wan C, Gu Y, Wang D, Yu H. Manganese promotes α-synuclein amyloid aggregation through the induction of protein phase transition. J Biol Chem. 2022;298:101469.

32. Huang S, Xu B, Liu Y. Calcium promotes α-synuclein liquid-liquid phase separation to accelerate amyloid aggregation. Biochem Biophys Res Commun. 2022;603:13–20.

33. Xu B, Chen J, Liu Y. Curcumin interacts with α-synuclein condensates to inhibit amyloid aggregation under phase separation. ACS Omega. 2022;7:30281–30290.

34. Burke KA, Janke AM, Rhine CL, Fawzi NL. Residue-by-residue view of in vitro FUS granules that bind the C-terminal domain of RNA polymerase II. Mol Cell. 2015;60:231–241.

35. Vernon RM, Chong PA, Tsang B, Kim TH, Bah A, Farber P, Lin H, Forman-Kay JD. Pi-Pi contacts are an overlooked protein feature relevant to phase separation. eLife. 2018;7:e31486.

36. Fung HYJ, Birol M, Rhoades E. IDPs in macromolecular complexes: the roles of multivalent interactions in diverse assemblies. Curr Opin Struct Biol. 2018;49:36–43.

37. Murthy AC, Dignon GL, Kan Y, Zerze GH, Parekh SH, Mittal J, Fawzi NL. Molecular interactions underlying liquid− liquid phase separation of the FUS low-complexity domain. Nat Struct Mol Biol. 2019;26:637–648.

38. Qamar S, Wang G, Randle SJ, Ruggeri FS, Varela JA, Lin JQ, Phillips EC, Miyashita A, Williams D, Ströhl F. FUS phase separation is modulated by a molecular chaperone and methylation of arginine cation-π interactions. Cell. 2018;173:720–734.

39. Li H-R, Chen T-C, Hsiao C-L, Shi L, Chou C-Y, Huang J-r. The physical forces mediating self-association and phase-separation in the C-terminal domain of TDP-43. Biochim Biophys Acta. 2018;1866:214–223.

40. Li H-R, Chiang W-C, Chou P-C, Wang W-J, Huang J-r. TAR DNA-binding protein 43 (TDP-43) liquid–liquid phase separation is mediated by just a few aromatic residues. J Biol Chem. 2018;293:6090–6098.

41. Schmidt HB, Barreau A, Rohatgi R. Phase separation-deficient TDP43 remains functional in splicing. Nat Commun. 2019;10:4890.

42. Letunic I, Bork P. 20 years of the SMART protein domain annotation resource. Nucleic Acids Res. 2018;46:D493–D496.

43. Wootton JC, Federhen S. Statistics of local complexity in amino acid sequences and sequence databases. Computers Chem. 1993;17:149–163.

44. Wootton JC. Sequences with ‘unusual’ amino acid compositions. Curr Opin Struct Biol. 1994;4:413–421.

45. Wootton JC, Federhen S. Analysis of compositionally biased regions in sequence databases. Methods Enzymol. 1996;266:554–571.

46. Kumar ST, Mahul-Mellier A-L, Hegde RN, Rivière G, Moons R, Ibáñez de Opakua A, Magalhães P, Rostami I, Donzelli S, Sobott F. A NAC domain mutation (E83Q) unlocks the pathogenicity of human alpha-synuclein and recapitulates its pathological diversity. Sci Adv. 2022;8:eabn0044.

47. Mehra S, Gadhe L, Bera R, Sawner AS, Maji SK. Structural and functional insights into αsynuclein fibril polymorphism. Biomolecules. 2021;11:1419.

48. Ptitsyn O. Molten globule and protein folding. Adv Protein Chem. 1995;47:83–229.

49. Deleage G, Roux B. An algorithm for protein secondary structure prediction based on class prediction. Protein Eng Des Sel. 1987;1:289–294.

50. Mészáros B, Erdős G, Dosztányi Z. IUPred2A: context-dependent prediction of protein disorder as a function of redox state and protein binding. Nucleic Acids Res. 2018;46:W329–W337.

51. Shaltiel-Karyo R, Frenkel-Pinter M, Rockenstein E, Patrick C, Levy-Sakin M, Schiller A, Egoz-Matia N, Masliah E, Segal D, Gazit E. A blood-brain barrier (BBB) disrupter is also a potent α-synuclein (α-syn) aggregation inhibitor: a novel dual mechanism of mannitol for the treatment of Parkinson disease (PD). J Biol Chem. 2013;288:17579–17588.

52. Taylor NO, Wei M-T, Stone HA, Brangwynne CP. Quantifying dynamics in phase-separated condensates using fluorescence recovery after photobleaching. Biophys J. 2019;117:1285–1300.

