## Supporting Information for "Liquid–liquid phase separation of α-synuclein is highly sensitive to sequence complexity"

**<u>Table of Contents</u>**

1. SUPPLEMENTARY FIGURES and TABLE
2. SUPPLEMENTARY TEXT

### 1. SUPPLEMENTARY FIGURES and TABLE

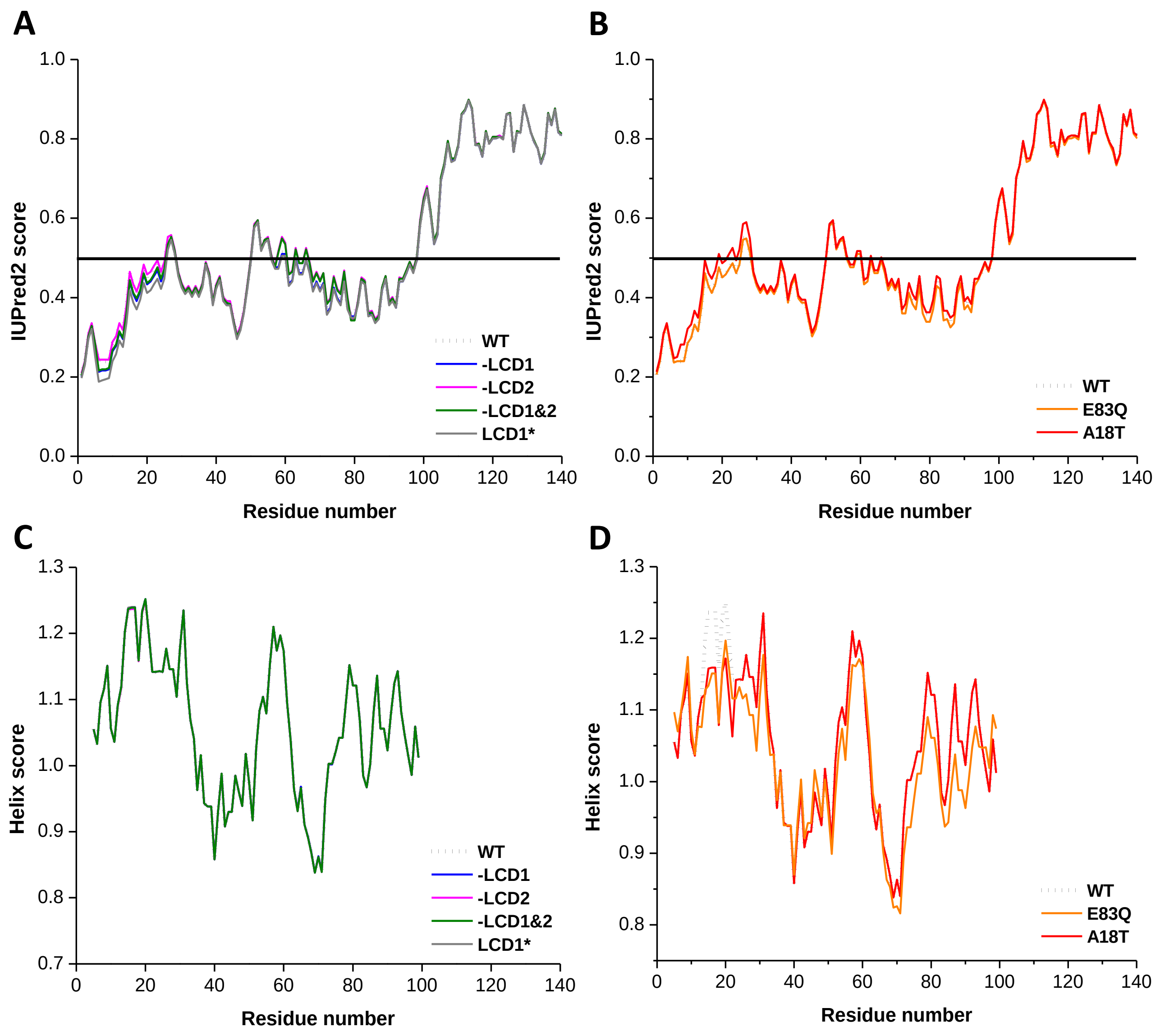


**Figure S1. (A)** and **(B)** Residue wise disorder scores of all designed and disease variants respectively, calculated using IUPred2A, showing lack of significant differences relative to WT in terms of disorder. **(C)** and **(D)** Residue wise α-helix scores of all designed and disease variants respectively, calculated using ProtScale by ExPASy, showing how each of the designed variants has exactly the same helix-forming propensity as the WT protein. However, patient mutations E83Q and A18T have slightly different helix-forming propensities.

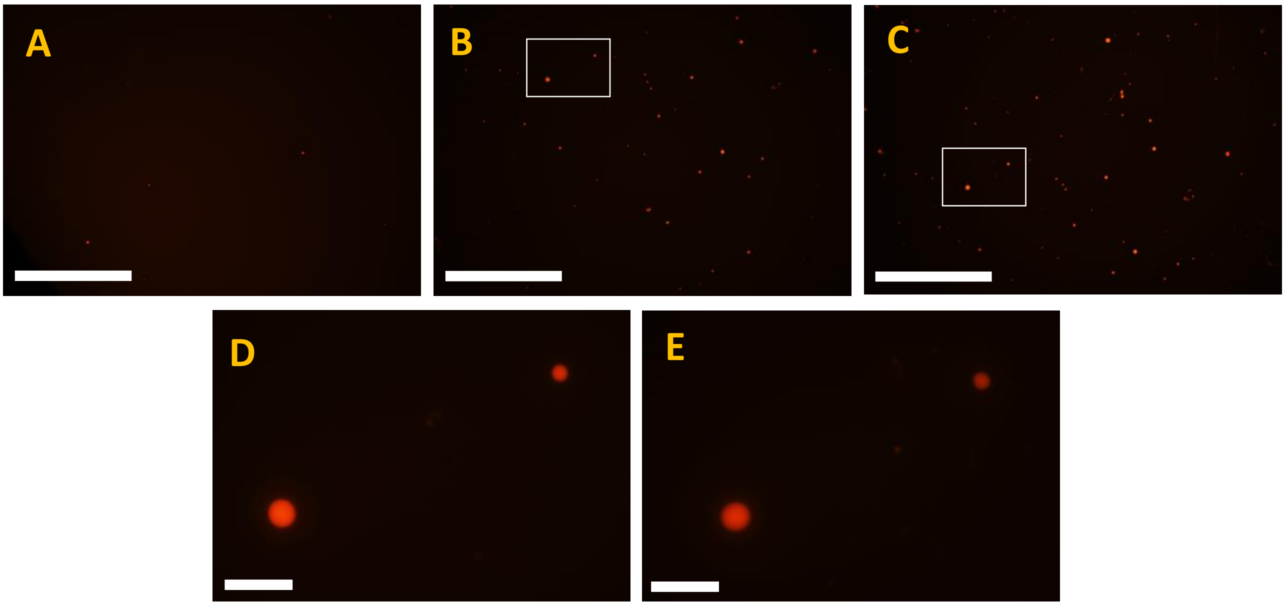


**Figure S2. (A)** to **(C)** Representative low magnification (10x) fluorescence microscopic images of a sample containing rhodamine-labelled WT α-syn in PBS incubated in presence of 20% PEG at 37 $^{0}$C for 0, 1 and 4 hours respectively, showing no visible increase in droplet size or number after 1 hour. Scale bar = 500 µm. **(D)** and **(E)** Higher magnification (60x) images of the same two droplets from the same sample at 1 and 4 hours respectively, showing no change in droplet size for either of them after 1 hour. Scale bar = 50 µm. Higher magnification image of the 0-hour sample could not be recorded as droplets at that stage were too mobile to be focussed on at 60x magnification.

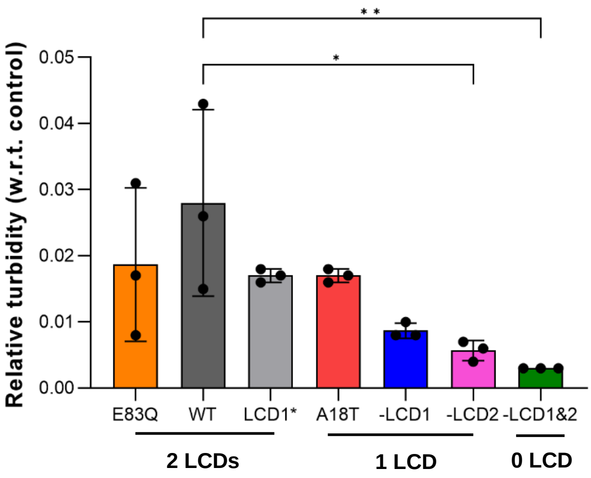


**Figure S3.** Turbidity ($OD_{500\ nm}$) values of LLPS variants at pH 7 after 4 hours of incubation, relative to background control (20% PEG in PBS). Data represent mean ± standard deviation (SD) for n = 3 independent experiments. Statistical significances are indicated by *: $p \leq 0.05$, **: $p \leq 0.01$, ***: $p \leq 0.001$, ****: $p \leq 0.0001$.

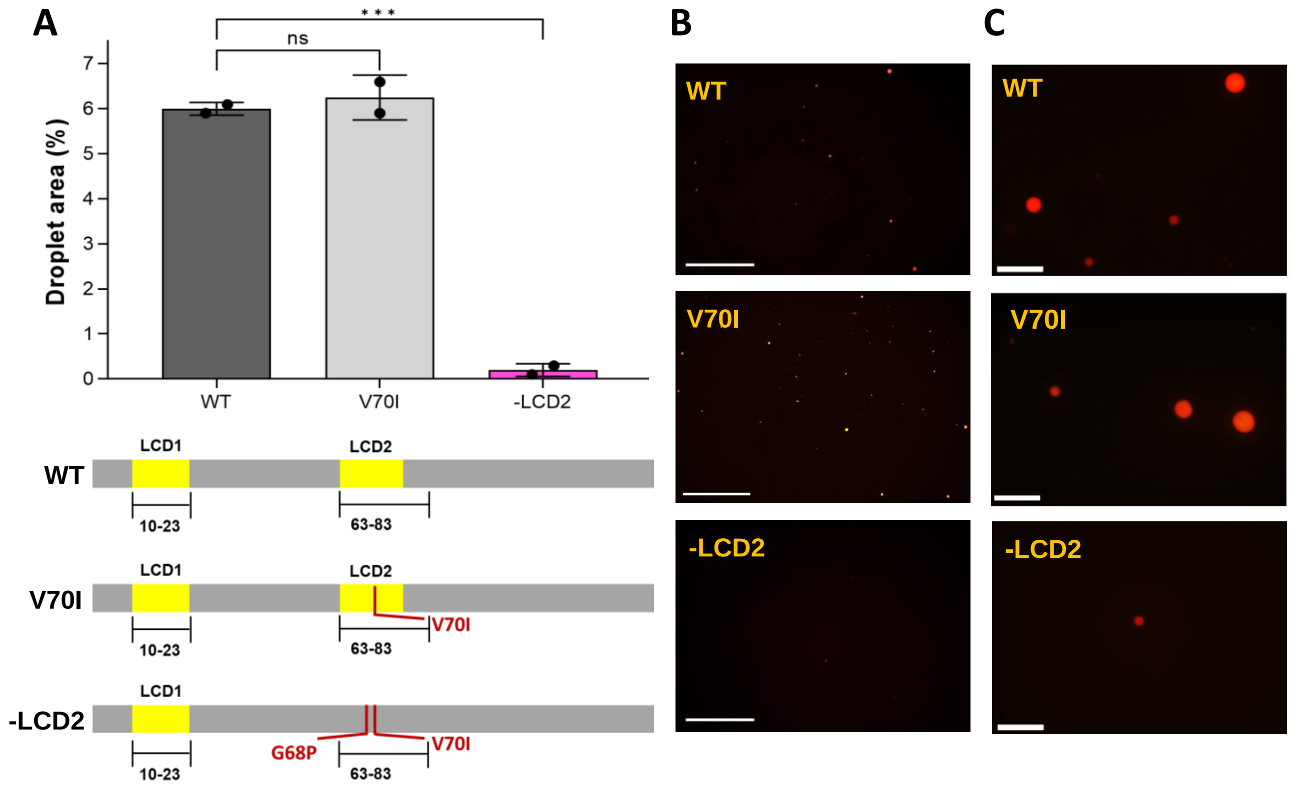


**Figure S4. (A)** *Top panel:* Quantification of droplet formation (from low magnification fluorescence microscopic images) for WT, V70I and –LCD2 variants of α-syn at pH 7. Data represent mean ± standard deviation (SD) for n = 2 independent experiments. Statistical significances are indicated by ns = non-significant, *: $p \leq 0.05$, **: $p \leq 0.01$, ***: $p \leq 0.001$, ****: $p \leq 0.0001$. *Bottom panel:* In silico analyses of amino acid sequences of WT, V70I and –LCD2 variants of α-syn using SMART, showing how the single V70I mutation does not disrupt LCD2 of the protein. **(B)** Representative low magnification (10x) fluorescence microscopic images of rhodamine-labelled WT, V70I, and -LCD2 variants respectively, incubated in PBS at pH 7, with 20% PEG, at 37 $^{0}$C for 4 hours. Scale bar = 500 µm. **(C)** Representative high magnification (60x) fluorescence microscopic images of rhodamine-labelled WT, V70I, and -LCD2 variants respectively, incubated in PBS at pH 7, with 20% PEG, at 37 $^{0}$C for 4 hours. Scale bar = 50 µm.

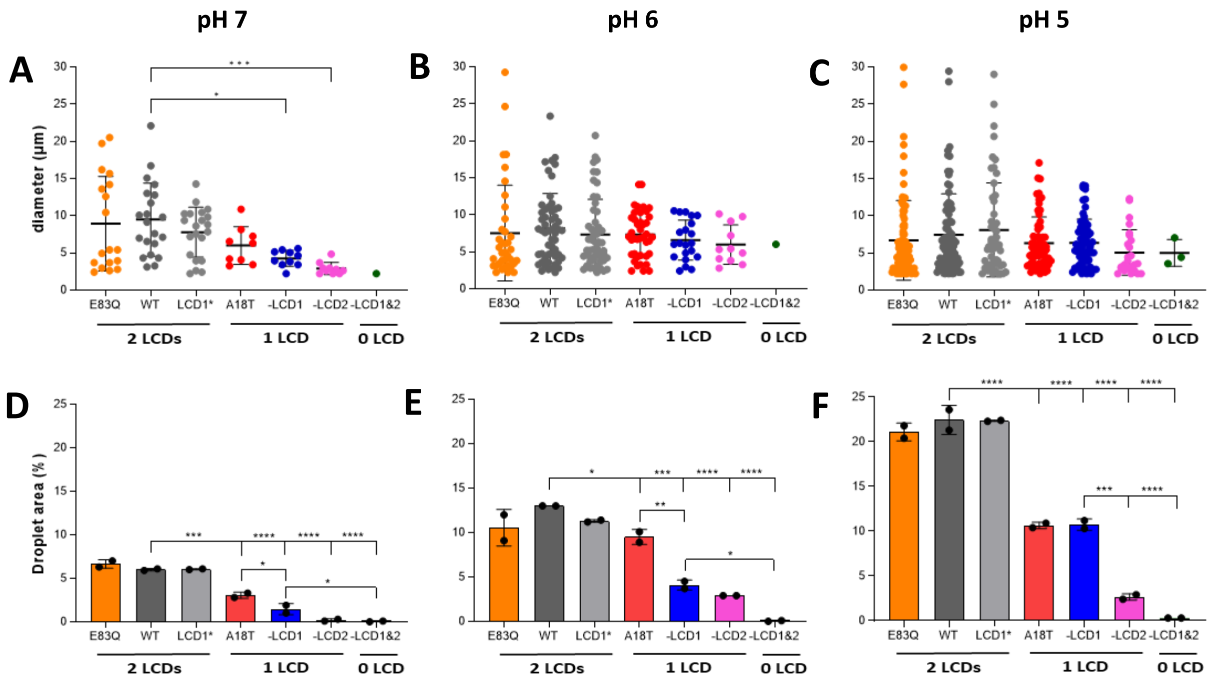


**Figure S5. (A)** to **(C)** Size (diameter) distribution analyses of droplets formed for all LLPS variants of α-syn, showing minimal differences in average size of droplets across variants and pH. However, number of droplets, and the maximum droplet diameter decreases consistently with decrease in number of LCDs across all pH, and lower pH leads to a greater number of droplets and higher values of the maximum size for all variants. **(D)** to **(F)** Quantification of area covered by droplets (average size x number) for all LLPS variants of α-Syn, showing significant reduction in droplet area with decreasing number of LCDs at all three pH. Data represent mean ± standard deviation (SD) for n = 2 independent experiments. Statistical significances are indicated by *: $p \leq 0.05$, **: $p \leq 0.01$, ***: $p \leq 0.001$, ****: $p \leq 0.0001$.

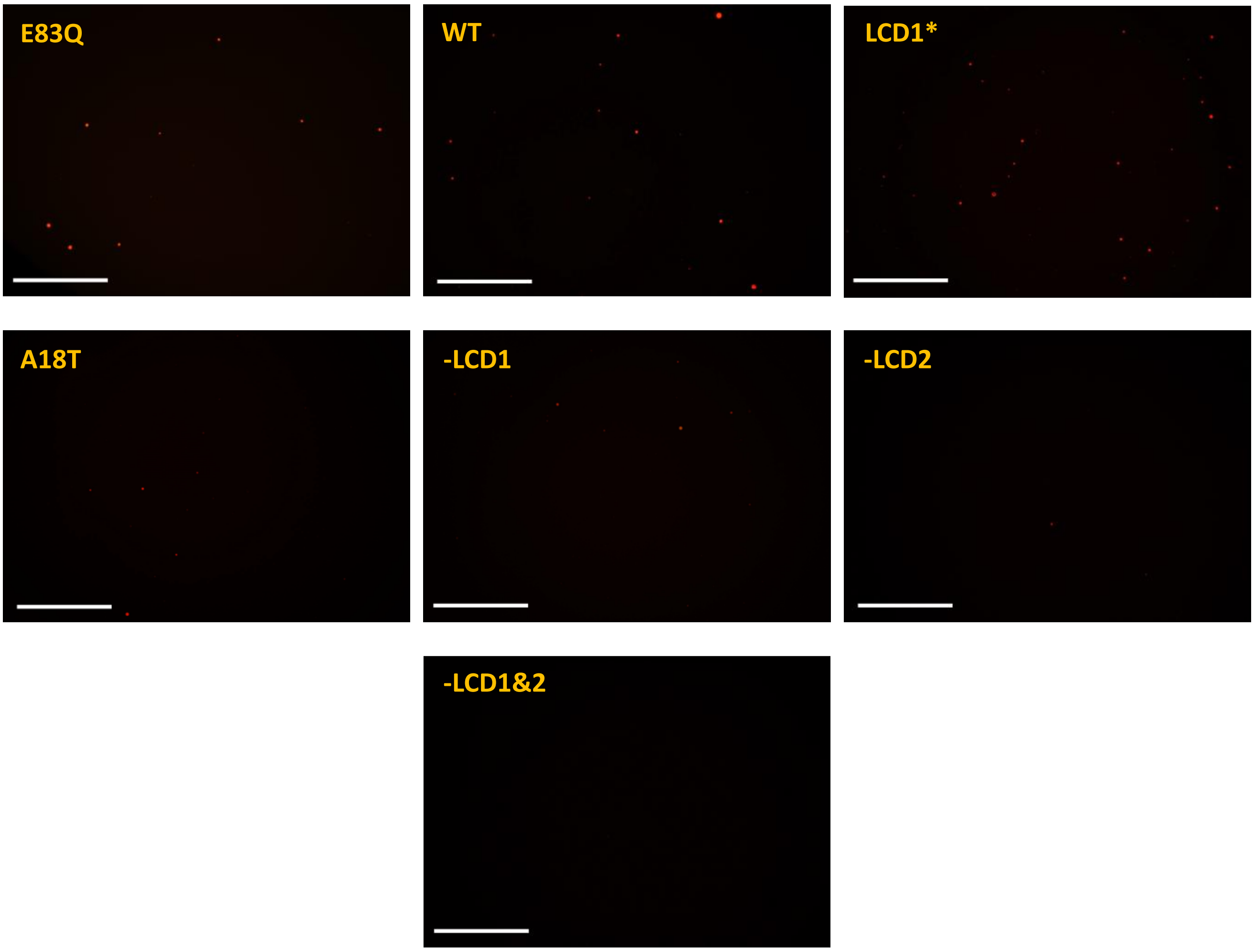


**Figure S6.** Representative low magnification (10x) fluorescence microscopic images of rhodamine-labelled α-syn variants incubated in PBS at pH 7, with 20% PEG, at 37 $^{0}$C for 4 hours. Scale bar = 500 µm.

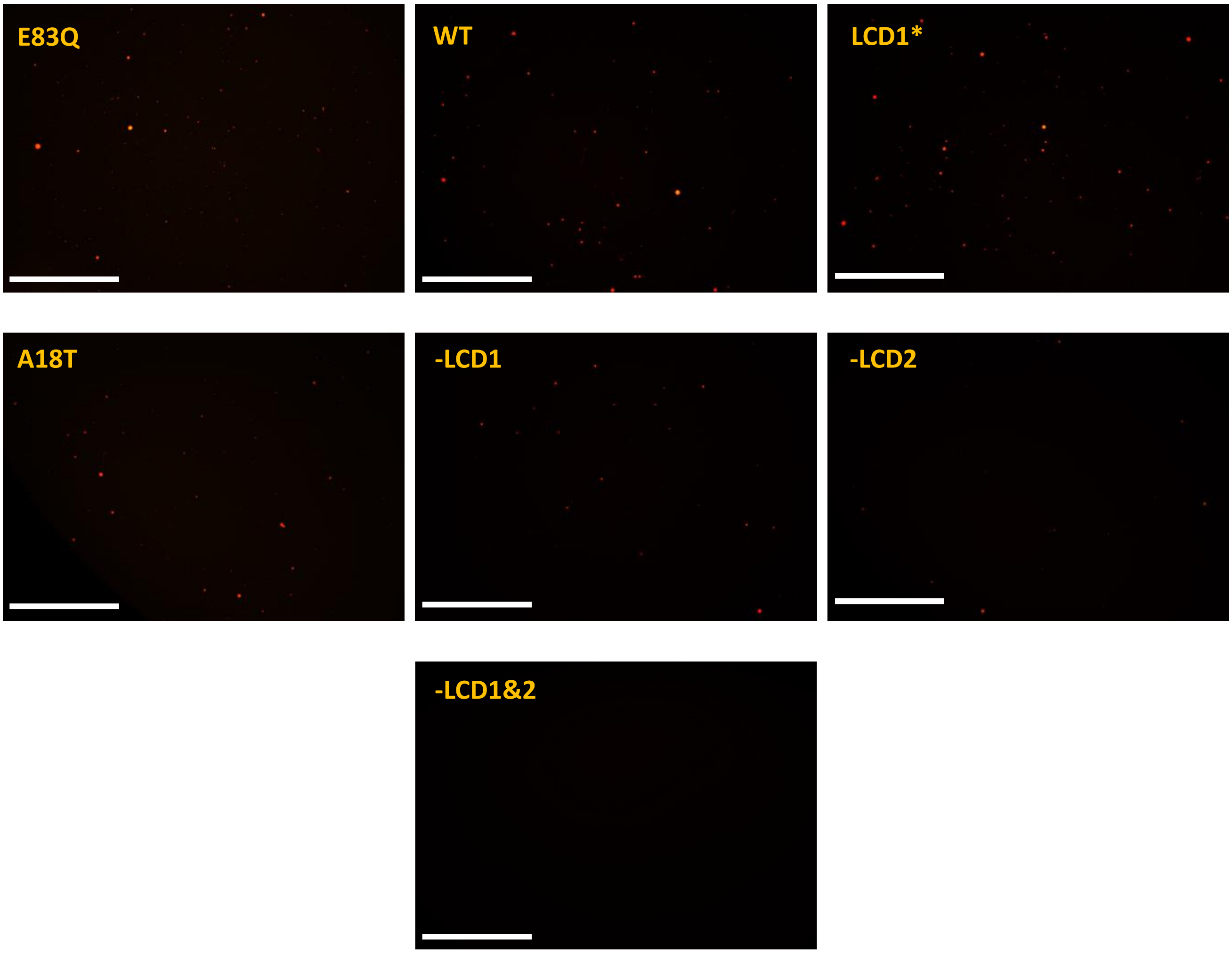


**Figure S7.** Representative low magnification (10x) fluorescence microscopic images of rhodamine-labelled α-syn variants incubated in PBS at pH 6, with 20% PEG, at 37 $^{0}$C for 4 hours. Scale bar = 500 µm.

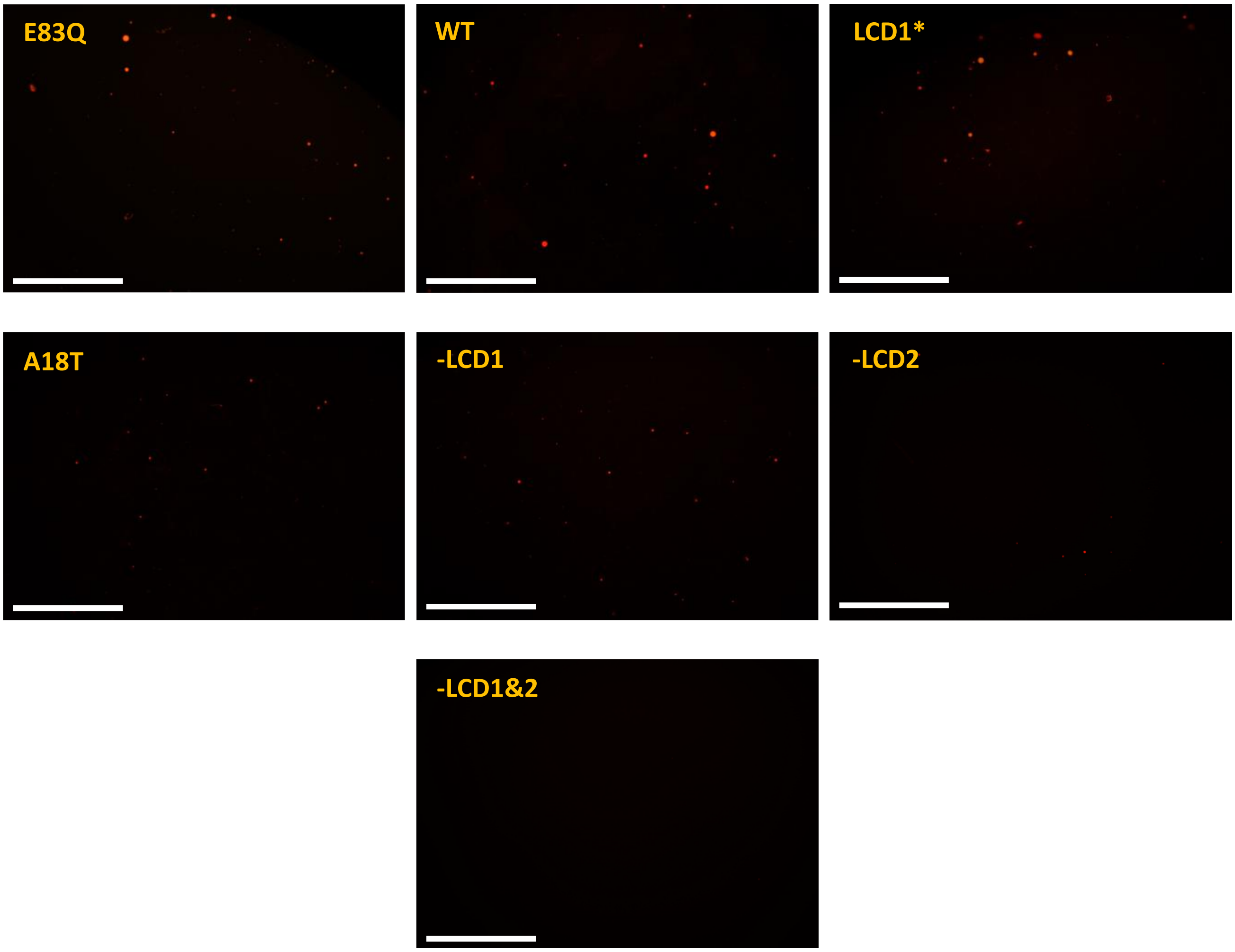


**Figure S8.** Representative low magnification (10x) fluorescence microscopic images of rhodamine-labelled α-syn variants incubated in PBS at pH 5, with 20% PEG, at 37 $^{0}$C for 4 hours. Scale bar = 500 µm.

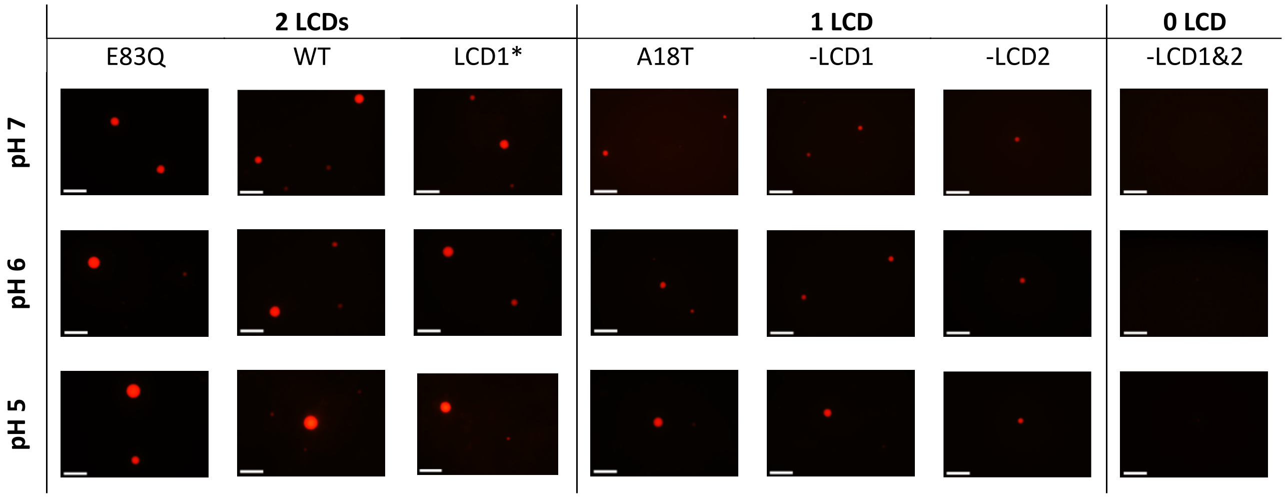


**Figure S9.** Representative high magnification (60x) fluorescence microscopic images of rhodamine-labelled LLPS variants of α-syn, incubated in PBS at pH 7, 6 and 5, with 20% PEG, at 37 $^{0}$C for 4 hours. Scale bar = 50 µm.

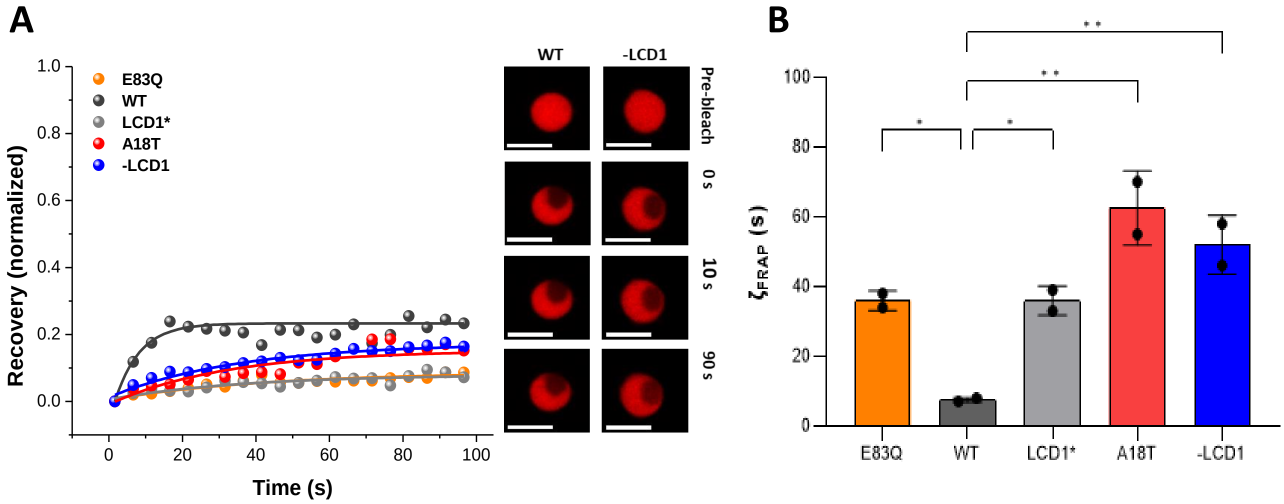


**Figure S10.** **(A)** *Left panel:* Normalized (and corrected for both passive bleaching and background) fluorescence recovery curves of different rhodamine-labelled LLPS variants (at 1 hour of incubation), showing a slower rate of recovery for all mutants compared to the WT protein. The recovery percentage is very low (<20%) for all the variants, indicating droplet maturation within an hour of formation. *Right panel:* Representative confocal images of WT and –LCD1 droplets at pre-bleach, bleach (0 s) and two post-bleach time points (10 and 90 s). Scale bar = 5 µm. Radius of the bleached region (ROI 1) = 2 µm. **(B)** FRAP recovery times of LLPS variants, showing the recovery times of the mutants to be significantly higher than that of the WT. Data represent mean ± standard deviation (SD) for n = 2 independent experiments. Statistical significances are indicated by *: $p \leq 0.05$, **: $p \leq 0.01$, ***: $p \leq 0.001$, ****: $p \leq 0.0001$. Mutants –LCD2 and –LCD1&2 could not be studied as the number and brightness of droplets in those samples were too low to focus on for photobleaching under our confocal microscope.

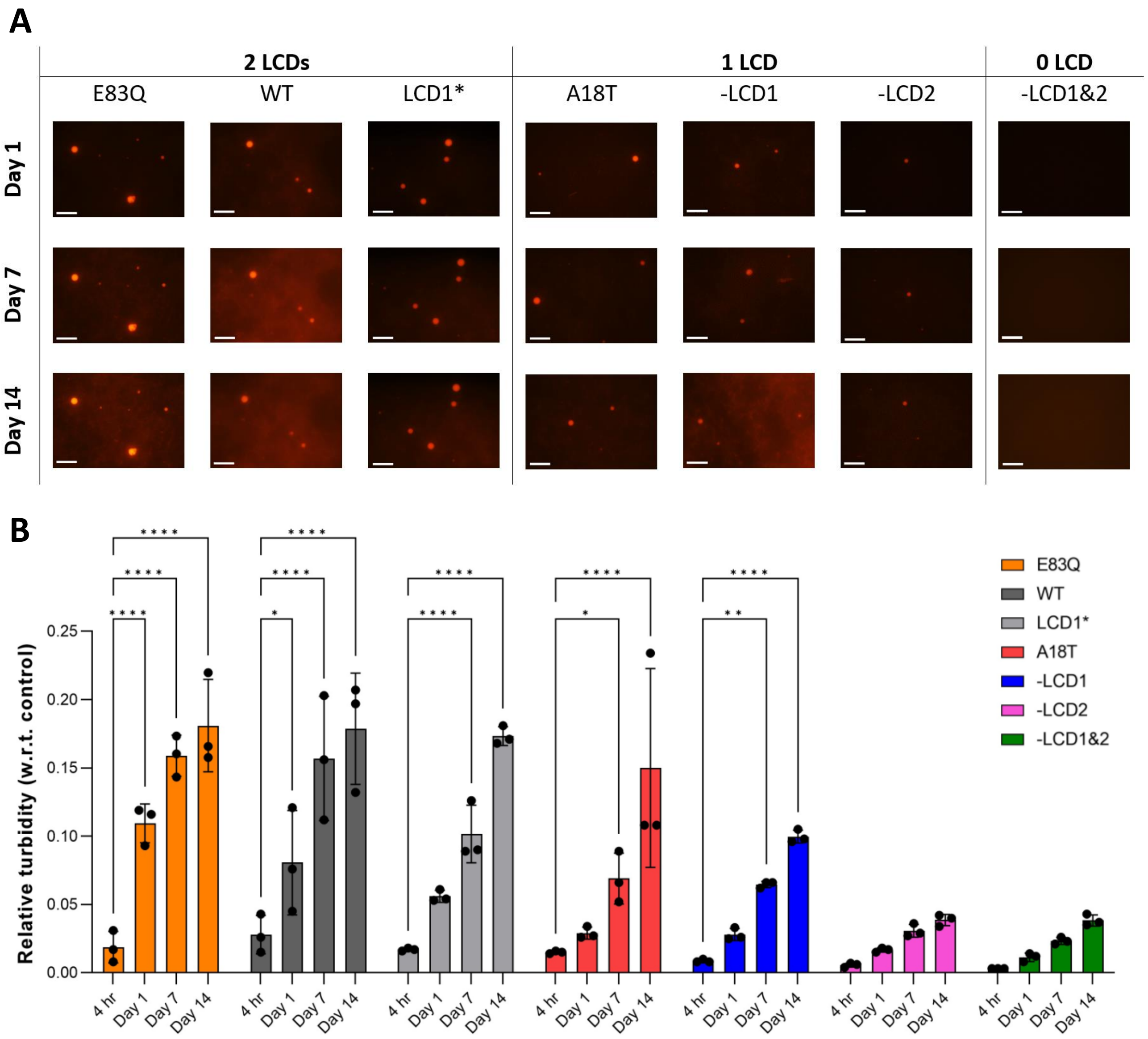


**Figure S11. (A)** Representative high magnification (60x) fluorescence microscopic images of rhodamine-labelled LLPS variants of α-Syn, in PBS at pH 7 with 20% PEG, after 1, 7 and 14 days of incubation at 37 $^{0}$C. Scale bar = 50 µm. **(B)** Turbidity ($OD_{500\ nm}$) values of all LLPS variants at pH 7 after 4 hours, day 1, day 7 and day 14 of incubation at 37 $^{0}$C, relative to background control (20% PEG in PBS) at each time point. Data represent mean ± standard deviation (SD) for n = 3 independent experiments. Statistical significances are indicated by *: $p \leq 0.05$, **: $p \leq 0.01$, ***: $p \leq 0.001$, ****: $p \leq 0.0001$.

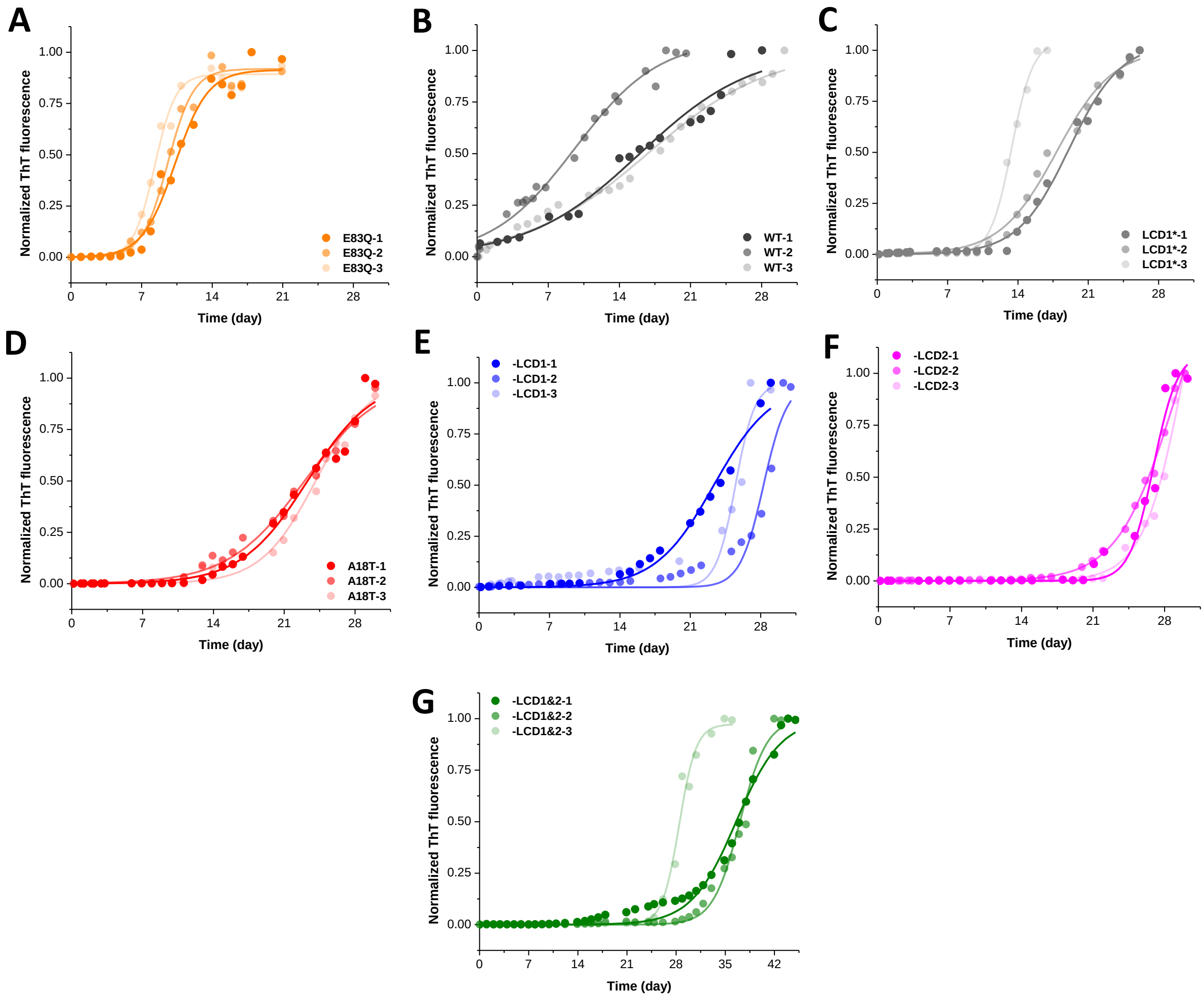


**Figure S12. (A)** to **(G)** Representative ThT-binding curves (normalized to saturation value) of E83Q, WT, LCD1*, A18T, -LCD1, -LCD2, and -LCD1&2 variants respectively, aggregated under LLPS conditions at pH 7.

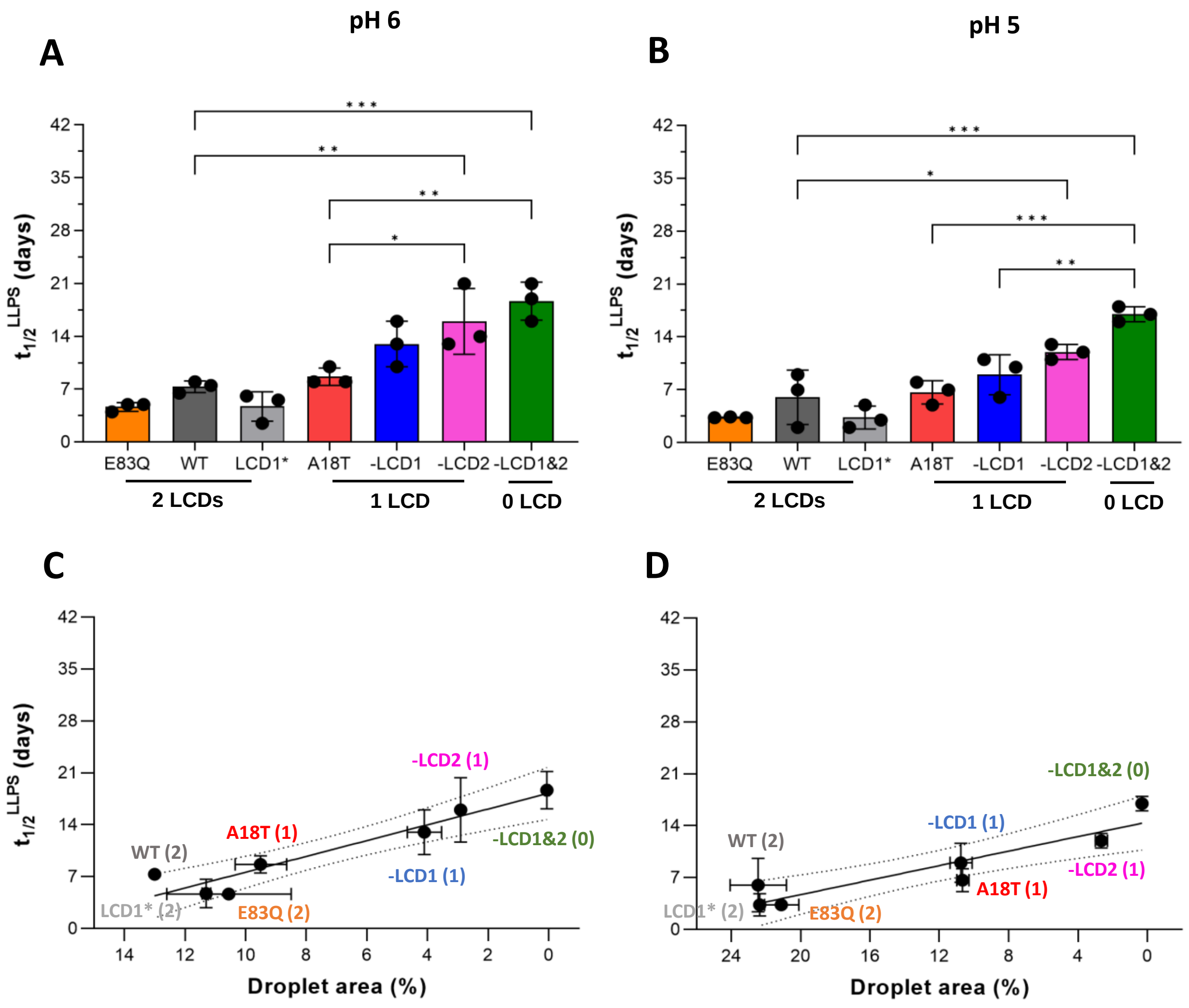


**Figure S13. (A)** and **(B)** Calculated half-times of fibrillation of all α-syn variants aggregated under LLPS conditions at pH 6 and 5 respectively. Data represent mean ± standard deviation (SD) for n = 3 independent experiments. Statistical significances are indicated by *: $p \leq 0.05$, **: $p \leq 0.01$, ***: $p \leq 0.001$, ****: $p \leq 0.0001$. **(C)** and **(D)** Correlation between half-times of fibrillation and LLPS propensities at pH 6 and 5 respectively. Numbers in brackets indicate number of LCDs. Dotted lines represent 95% confidence interval for linear regression.

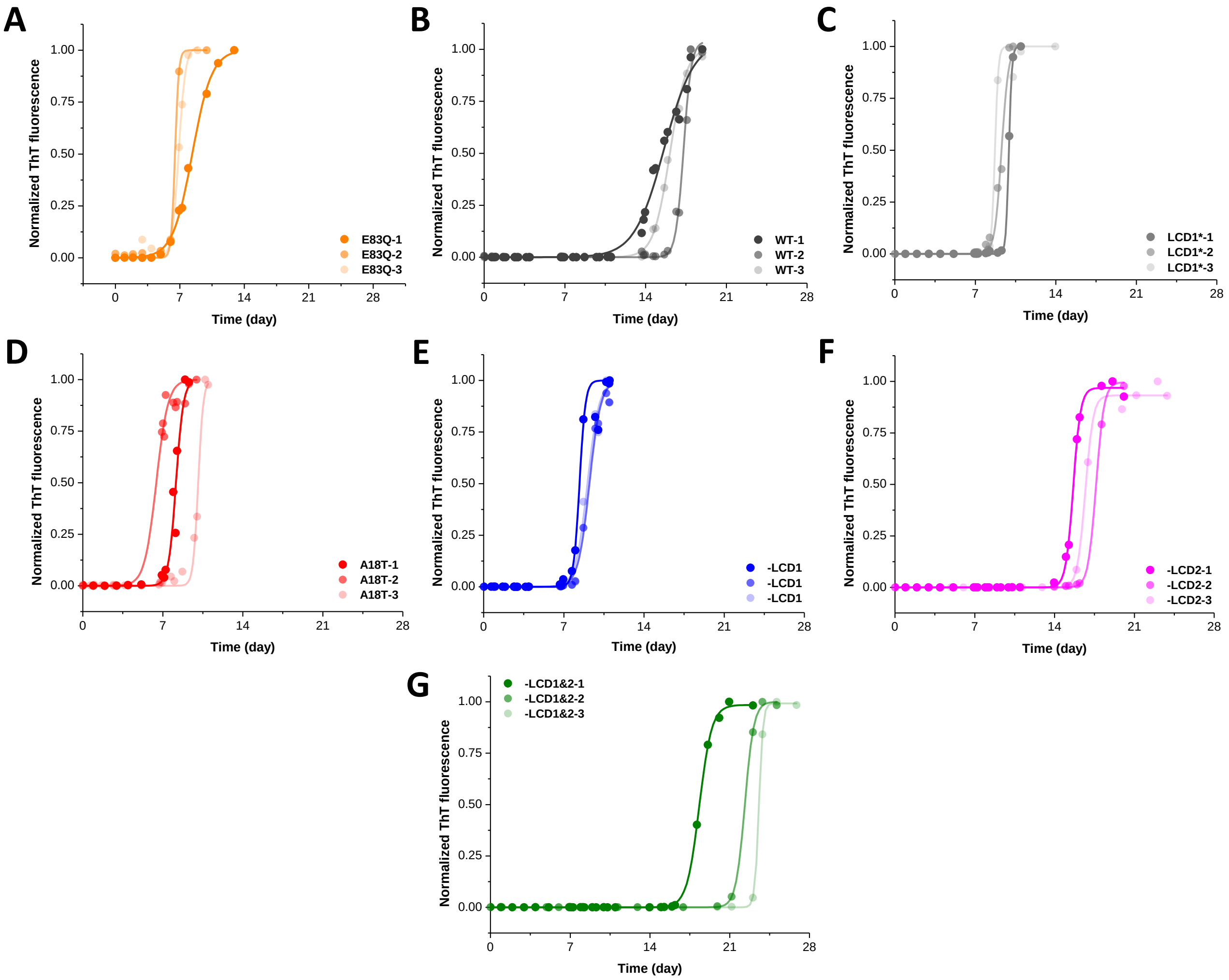


**Figure S14. (A)** to **(G)** Representative ThT-binding curves (normalized to saturation value) of E83Q, WT, LCD1*, A18T, -LCD1, -LCD2, and -LCD1&2 variants respectively, aggregated under non-LLPS conditions at pH 7.

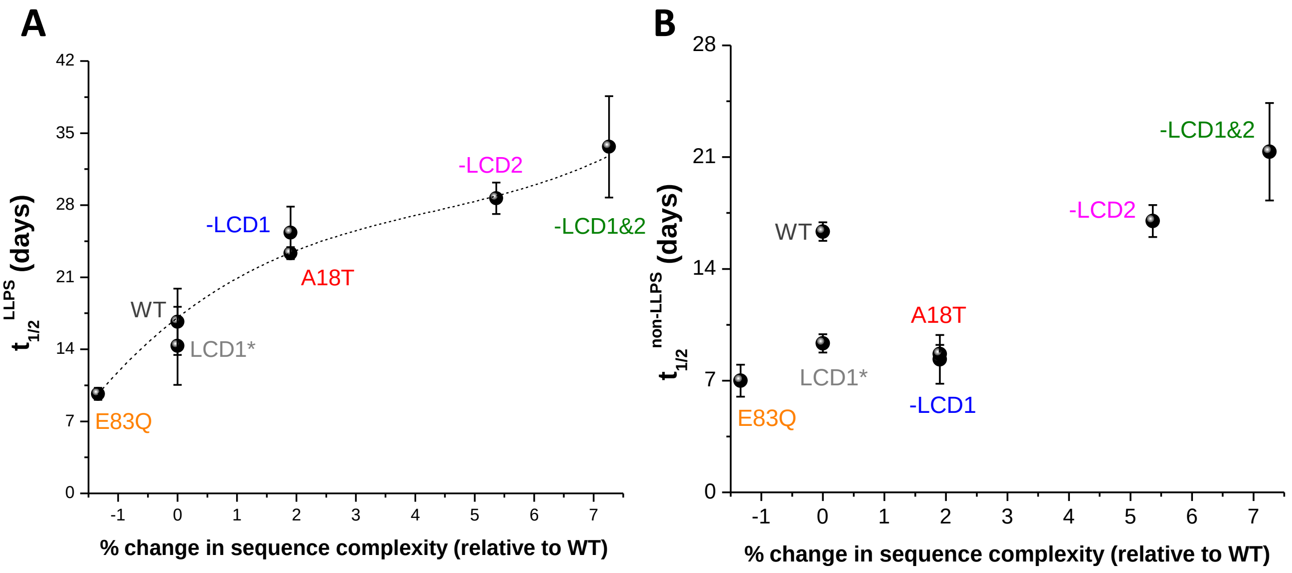


**Figure S15. (A)** Fibrillation half-times of α-syn variants aggregated under LLPS conditions increase with increasing sequence complexity. **(B)** Fibrillation half-times of α-syn variants aggregated under non-LLPS conditions do not correlate with sequence complexity. Data represent mean ± standard deviation (SD) for n = 3 independent experiments.

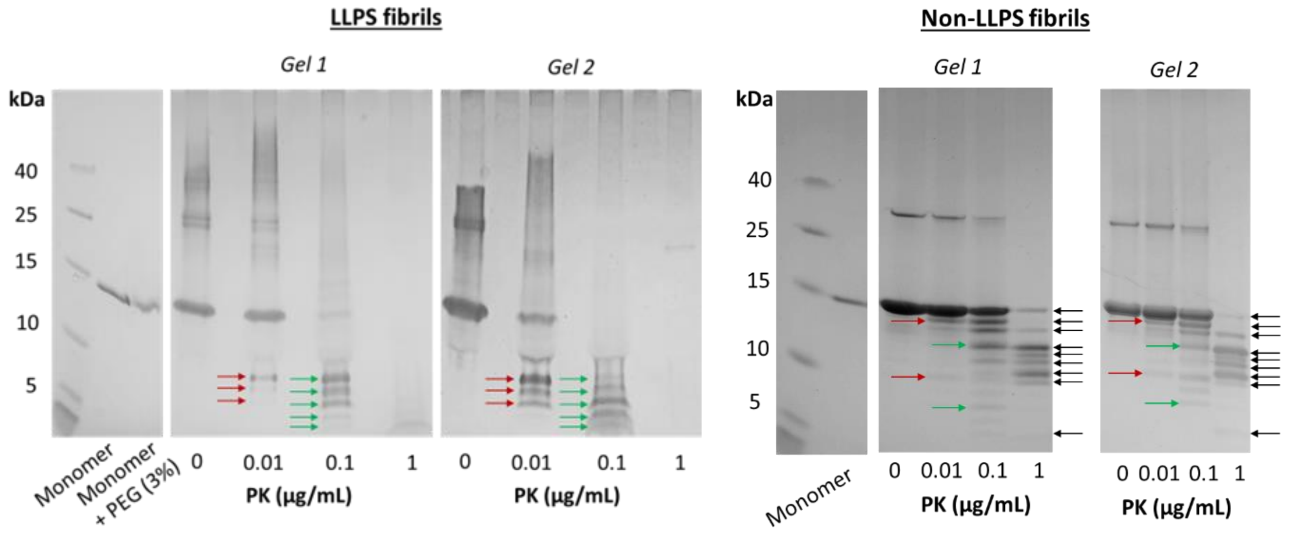


**Figure S16.** Gel images corresponding to two independent limited proteolysis experiments with WT α-syn fibrils produced under LLPS and non-LLPS conditions, showing robust fingerprinting of each fibril type.

**Table S1.** Calculated hydrophobicity scores, charges at pH 7.4, and isoelectric points (pI) of designed and disease (italicized) variants of α-syn.

| **Mutation(s)** | **Variant Name** | **GRAVY*** | **Charge at pH 7.4** | **pI** |
|---|---|---|---|---|
| None | WT | -0.403 | -9.7 | 4.67 |
| V16I | -LCD1 | -0.400 | -9.7 | 4.67 |
| G68P, V70I | -LCD2 | -0.409 | -9.7 | 4.67 |
| V16I, G68P, V70I | -LCD1&2 | -0.407 | -9.7 | 4.67 |
| V15I, V16I | LCD1* | -0.399 | -9.7 | 4.67 |
| *E83Q* | *E83Q* | *-0.403* | *-8.7* | *4.73* |
| *A18T* | *A18T* | *-0.421* | *-9.7* | *4.67* |

*The grand average of hydropathy (**GRAVY**) value for protein sequences. It is defined by the sum of hydropathy values of all amino acids divided by the protein length.

#### 2. SUPPLEMENTARY TEXT:

**Fluorescence recovery after photobleaching (FRAP) experiments on phase-separated α-syn variants:** In the presence of PEG 8000 in PBS, droplets of WT α-syn started forming almost immediately (Fig. S2A), and upon continued incubation at 37 $^{0}$C, they matured within an hour (Fig. S2B). When we performed FRAP experiments on the 1-hour droplets of all variants, their observed low fluorescence recovery percentages confirmed that droplets formed under our experimental conditions became solid-like within an hour of their formation (Fig. S10A). Upon analysis of the FRAP recovery times ($\tau_{FRAP}$) of all variants, we found that all mutants took significantly higher time to recover relative to the WT (Fig. S10B), indicating reduced dynamicity within droplets. Therefore, although the extent of droplet formation of α-syn positively correlated to its sequence complexity, its dynamicity within the droplets did not.
